# Nucleocytoplasmic Proteomic Analysis Uncovers eRF1 and Nonsense Mediated Decay as Modifiers of ALS *C9orf72* Toxicity

**DOI:** 10.1101/677419

**Authors:** Juan A. Ortega, Elizabeth L. Daley, Sukhleen Kour, Marisa Samani, Liana Tellez, Haley S. Smith, Elizabeth A. Hall, Y. Taylan Esengul, Yung-Hsu Tsai, Tania F. Gendron, Christopher J. Donnelly, Teepu Siddique, Jeffrey N. Savas, Udai B. Pandey, Evangelos Kiskinis

## Abstract

The most common genetic cause of amyotrophic lateral sclerosis (ALS) and frontotemporal dementia (FTD) is a hexanucleotide repeat expansion in *C9orf72* (C9-HRE). While RNA and dipeptide repeats produced by the C9-HRE disrupt nucleocytoplasmic transport, the proteins that become redistributed remain unknown. Here, we utilized subcellular fractionation coupled with tandem mass spectrometry and identified 126 proteins, enriched for protein translation and RNA metabolism pathways, which collectively drive a shift towards a more cytosolic proteome in C9-HRE cells. Amongst these was eRF1, which regulates translation termination and nonsense-mediated decay (NMD). eRF1 accumulates within elaborate nuclear envelope invaginations in patient iPSC-neurons and postmortem tissue and mediates a protective shift from protein translation to NMD-dependent mRNA degradation. Overexpression of eRF1 and the NMD-driver UPF1 ameliorate C9-HRE toxicity *in vivo*. Our findings provide a resource for proteome-wide nucleocytoplasmic alterations across neurodegeneration-associated repeat expansion mutations and highlight eRF1 and NMD as therapeutic targets in *C9orf72*-associated ALS/FTD.

## INTRODUCTION

A hexanucleotide repeat expansion (HRE) in the first intron of *C9orf72* is the most common genetic cause of ALS and FTD (Taylor et al., 2016; Vucic et al., 2014). It accounts for as much as 25% and 40% of familial ALS and FTD respectively, and 5-7% of all sporadic ALS cases. Healthy individuals typically carry less than 30 GGGGCC (G_4_C_2_) repeats, while ALS and FTD patients carry hundreds or thousands (DeJesus-Hernandez et al., 2011; Renton et al., 2011). The *C9orf72* HRE (C9-HRE) is proposed to drive neurodegeneration through three non-exclusive mechanisms (Balendra and Isaacs, 2018; Gendron et al., 2014; Gitler and Tsuiji, 2016; Haeusler et al., 2016). *C9orf72* expression is lower in patients suggesting that haploinsufficiency contributes to pathogenesis (Shi et al., 2018; van Blitterswijk et al., 2015). Gain-of-function neurotoxic effects occur through production of aberrant C9-HRE RNA (Donnelly et al., 2013), and through dipeptide repeat proteins (DPRs) translated by a non-ATG dependent mechanism from the HRE (Mori et al., 2013a; Zu et al., 2013). The G_4_C_2_ repeat is bi-directionally transcribed and corresponding sense and antisense RNA accumulates in predominantly nuclear RNA foci in patient cells and tissue (Cooper-Knock et al., 2015; Donnelly et al., 2013; Zu et al., 2013). Moreover, five C9-HRE DPRs (GA, GP, GR, PR, PA) are found in patient cortex and spinal cord tissue (Gomez-Deza et al., 2015; Mackenzie et al., 2015).

While the relative contribution of loss-of-function effects and each gain-of-function effect (RNA or DPRs) remains unresolved, converging evidence suggests that the C9-HRE disrupts the balance of proteins transported between the nucleus and cytoplasm. The expanded C9-RNA binds and sequesters RNA-binding proteins in the nucleus (Donnelly et al., 2013). The arginine-rich DPRs bind and sequester many proteins with low complexity domains that assemble into membrane-less organelles through liquid-liquid phase separation (Lee et al., 2016; Lin et al., 2016). These include RNA granules, nucleoli and various components of the nuclear pore complex (NPC). Moreover, a series of genetic studies in C9-HRE *Drosophila* and yeast model systems identified nucleocytoplasmic (N/C) trafficking as a critical mediator of C9-HRE toxicity (Freibaum et al., 2015; Jovicic et al., 2015; Zhang et al., 2015). Specifically, NPCs and proteins involved in nuclear import and export are strong modifiers of C9-HRE toxicity. Importantly, the nature of the proteins that become redistributed or mislocalized due to these mechanisms remains unknown.

Abnormal N/C localization of proteins could also have broader relevance for sporadic disease as cytoplasmic accumulation of TDP-43, a protein typically found in the nucleus, is a neuropathological hallmark in most ALS patients (Neumann et al., 2006). Recent studies suggest that cytoplasmic aggregation of TDP-43 interferes with N/C transport, and TDP-43 protein that aggregates in the cytoplasm, specifically outside RNA-rich structures (Mann et al., 2019), is more toxic than TDP-43 that aggregates in the nucleus (Barmada et al., 2014; Woerner et al., 2016). At the same time, NPCs exhibit an age-dependent deterioration in postmitotic neurons, highlighting a potential interaction between mutations, such as the C9-HRE, and age-dependent neurodegenerative NPC dysfunction (D’Angelo et al., 2009; Savas et al., 2012; Toyama et al., 2013).

In this study, we sought to identify proteins with altered N/C distribution in C9-HRE carrier cells. We reasoned that proteins can become mislocalized due to defects in N/C transport, pathogenic sequestration within C9-RNA foci and C9-DPR aggregates, or redistribution to counteract the cellular defects driven by the mutation. We hypothesized that these alterations would highlight disease-relevant mechanisms. To test this hypothesis, we developed an unbiased approach to interrogate proteome-wide N/C protein distribution, and determined how it was affected by the C9-HRE. We identified a number of proteins that are redistributed in C9-HRE cells, primarily implicated in RNA metabolism and protein synthesis. Amongst the proteins that become more nuclear was eRF1, which regulates the balance between protein translation termination and nonsense mediated decay (NMD)-based mRNA degradation. In response to the C9-HRE, eRF1 triggers a protective shift from protein translation to UPF1-dependent NMD, and targets the expanded C9-HRE transcript for degradation. Our work strengthens the hypothesis that a major component of mutant *C9orf72* toxicity is related to mRNA processing defects, provides a link between the C9-HRE and the NMD pathway, and highlights eRF1 and UPF1 as novel therapeutic targets in *C9orf72*-associated ALS/FTD.

## RESULTS

### The *C9orf72* Expansion Leads to Proteome-Wide Nucleocytoplasmic Redistribution

To determine if the C9-HRE affects the N/C distribution of proteins, we expressed a control or disease-associated number of G_4_C_2_ repeat sequences (8 or 58 respectively) in HEK-293 cells (Figure 1A) (Freibaum et al., 2015). The constructs included a canonical start site (AUG) in frame with poly-GP DPR and GFP (Figures 1 and S1A). Poly-GP (AUG-driven) and poly-GR (non-AUG-driven) were both produced in transfected cells (Figure S1B-D), and the size of the G_4_C_2_-GFP fusion protein corresponded to the respective number of repeats by Western blot (WB) analysis (Figure S1E). We used fluorescent activated cell sorting (FACS) to purify transfected HEK-293 cells (Figures 1A and S1F-H), performed subcellular fractionation to isolate nuclear (N) and cytosolic (C) proteins, and analyzed the proteome by semi-quantitative mass spectrometry (MS) (Figure 1A). We corroborated the precision of the subcellular fractionation by: a) WB analysis that showed the nuclear marker TAFII and cytoplasmic marker VINCULIN confined to the correct fraction (Figure 1B); and b) localization analysis of well-characterized nuclear (n=37) and cytosolic (n=68) resident proteins by MS (Figure 1C and Table S1). C9×58 expression was not toxic at 48h, so we used this time point for MS analyses (Figures S1I-K).

**Figure 1.**
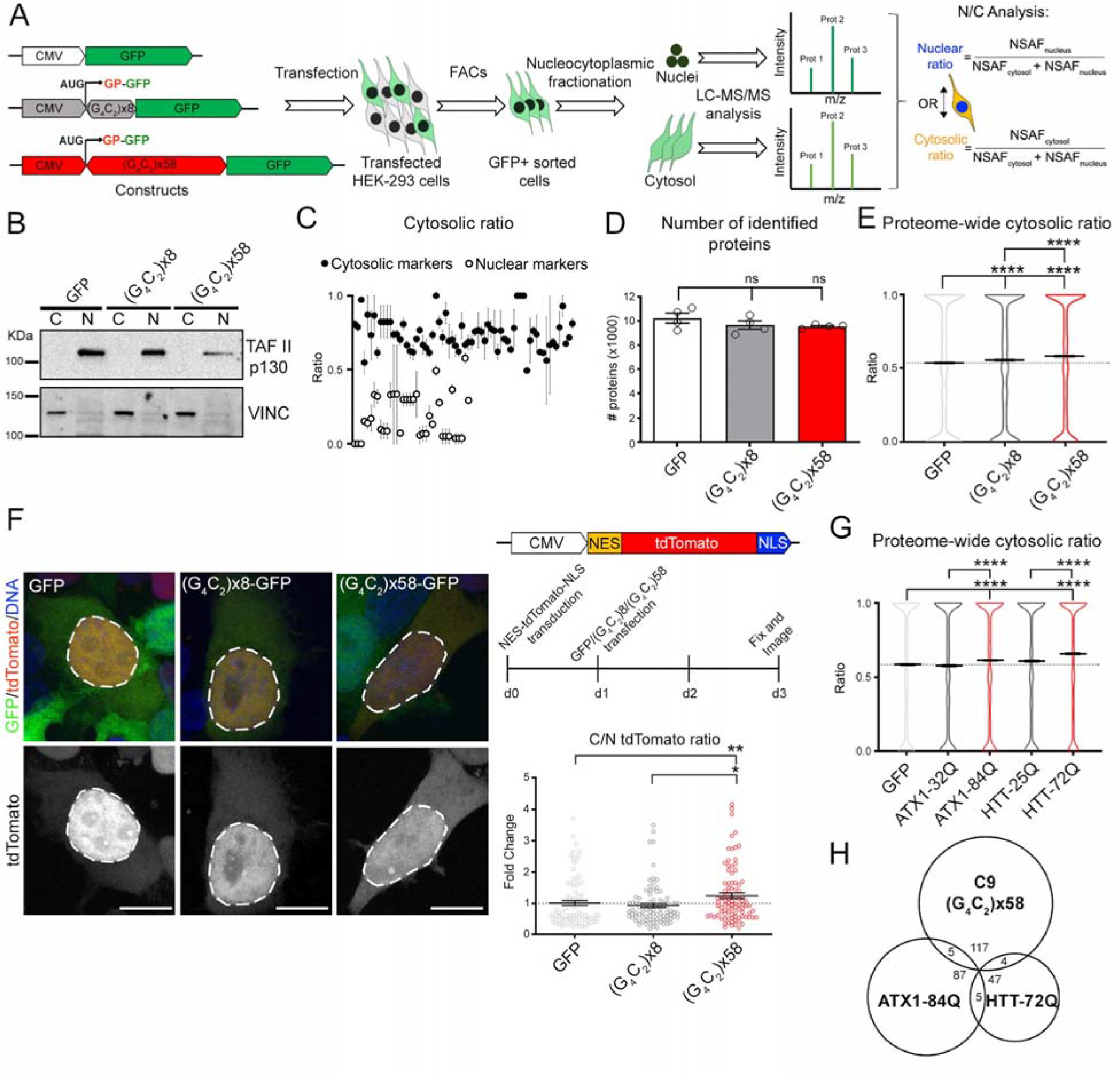
The C9-HRE Leads to Proteome-Wide N/C Redistribution. (A) Experimental schematic used to identify the effects of the C9-HRE on the subcellular proteome. (B) WB for TAFII p130 and VINCULIN in the cytosolic (C) and nuclear (N) fractions of GFP-, C9×8- and C9×58-expressing cells. (C) Dot plots representing the cytosolic ratio obtained from LC-MS/MS analysis in GFP-expressing cells of *bona fide* cytosolic (black) and nuclear (gray) markers (n=3 independent biological replicates). (D) Bar plots representing the average number of redundant proteins identified in cells expressing GFP, C9×8 and C9×58 by LC-MS/MS. ANOVA; ns=not significant. (E) Violin plots showing the proteome-wide cytosolic ratio of the ∼10,000 proteins identified in cells expressing GFP (0.528±0.003; mean ± SEM), C9×8 (0.548±0.003), and C9×58 (0.575±0.003). Kruskal-Wallis test. (F) Left: Representative confocal images of cells expressing N/C shuttling tdTomato reporter and GFP, C9×8-GFP or C9×58-GFP. Dashed lines mark the nuclear boundary. Right-top: N/C shuttling tdTomato reporter and workflow schematics. Right-bottom: Dot plot showing the fold change in the cytosolic/nuclear (C/N) ratio of the tdTomato signal in C9×58-expressing cells compared to controls. Individual cells are displayed as dots. Kruskal-Wallis test. (G) Violin plots showing the proteome-wide cytosolic ratio of the ∼8,000 proteins identified in cells expressing GFP (0.58±0.004; mean ± SEM), ATX1-32Q (0.57±0.004), ATX1-84Q (0.61±0.004), HTT-25Q (0.60±0.004) and HTT-72Q (0.65±0.004). Kruskal-Wallis test. (H) Venn diagram showing the level of overlap of redistributed proteins in cells expressing C9×58, ATX1-84Q and HTT-72Q. In graphs, bars and horizontal lines represent mean ± SEM, and dotted lines mark the GFP-transfected cells mean value. Scale bars: 15 μm.

We identified approximately 10,000 redundant proteins in each sample (GFP, C9×8, C9×58), with no statistically significant differences in the average protein number (n=4 independent biological replicates, m=8 of N and C fractions per sample, p=0.116) (Figure 1D). To assess subcellular protein distribution, we calculated the average N/C ratio for all proteins identified across experiments in the three samples. We then asked if expression of the C9-HRE affected the proteome-wide N/C distribution. We found the average cytosolic ratio of the proteome was significantly shifted to a higher level in the presence of 58 repeats, compared to 8 repeats or control GFP (n=4 independent biological replicates; GFP *vs*. C9×58: p<0.0001 and C9×8 *vs*. C9×58: p<0.0001) (Figures 1E and S1L). To investigate the effects on protein distribution by an alternative method, we used a tdTomato reporter containing a nuclear localization signal (NLS) and a nuclear export signal (NES) (Figure 1F, top right). We first validated that the reporter became even more nuclear (2-fold higher) after treatment with nuclear export inhibitor KPT-330 (Figure S1M). We then found that upon C9×58 expression the tdTomato signal became significantly more cytosolic relative to both controls (n=3 independent biological replicates; GFP *vs*. C9×58: p<0.01, 23% increase ±8; and C9×8 *vs*. C9×58: p=0.03, 36% increase ±7) (Figure 1F, left and bottom right). These results suggest that C9×58-expressing cells undergo global protein redistribution, with more proteins accumulating in the cytoplasm.

### Repeat Expansion Mutations Lead to Proteome-Wide Nucleocytoplasmic Redistribution of Distinct Proteins

We next asked if other neurodegeneration-associated repeat expansion mutations also affect the N/C distribution of proteins. To address this question, we used constructs expressing a polyglutamine (CAG, polyQ) expansion in the Ataxin 1 gene (*ATX1*-84Q), or in the first exon of the Huntingtin gene (*HTT*-72Q), that cause spinocerebellar ataxia type 1 (SCA1) and Huntington’s disease (HD), respectively. As controls, we used corresponding constructs with a poly-Q expansion below the threshold of disease (32Q and 25Q, respectively), and a GFP control vector (Figure S1N). Following FACS purification, subcellular fractionation and MS analysis (Figures S1N-Q), we found that expression of poly-Q repeats also caused a significant shift towards a more cytosolic proteome relative to controls (n=3 independent biological replicates, m=6 of N and C fractions per sample, p < 0.0001) (Figure 1G).

We next examined the nature of the proteins that become significantly redistributed and found very little overlap amongst all three repeat expansions, where only 4% and 3.2% of the proteins that shifted in C9×58-expressing cells were also affected by polyQ-ATX1 or HTT, respectively (Figure 1H and Table S1). Moreover, although both *ATX1* and *HTT* repeats represent polyQ sequences, the degree of overlap in redistributed proteins was similarly small (5.2% and 8.9%) (Figures 1H and Table S1). The reason for the differential effects could be associated with the biophysical or functional properties of each mutation, the subcellular localization of the mutant protein, or the size of the repeat itself. These results suggest that impaired N/C trafficking could be broadly implicated in other neurodegenerative diseases, corroborating recent studies for HD and Alzheimer disease models (Eftekharzadeh et al., 2018; Gasset-Rosa et al., 2017; Grima et al., 2017; Woerner et al., 2016).

### The *C9orf72* Repeat Expansion Causes a Redistribution of Proteins Involved in RNA Metabolism and Protein Translation

Comprehensive statistical analysis identified 126 unique proteins with a significantly altered N/C ratio in C9×58-expressing cells (n=4 independent biological replicates; Kruskal–Wallis rank test; C9×58 *vs*. GFP and C9×8: p<0.05) (Figure 2A and Table S1). These 126 proteins represented 1.2% of the total number of proteins identified and all showed a bidirectional change, with 56% shifting from the nucleus to cytosol and 44% shifting in the opposite direction (Figure 2B). Approximately 60% of proteins that relocated to the cytosol are normally nuclear, whereas 46% of proteins that relocated to the nucleus, are normally cytosolic (Figure S2A) (Binder et al., 2014), suggesting these changes may have critical functional ramifications.

**Figure 2.**
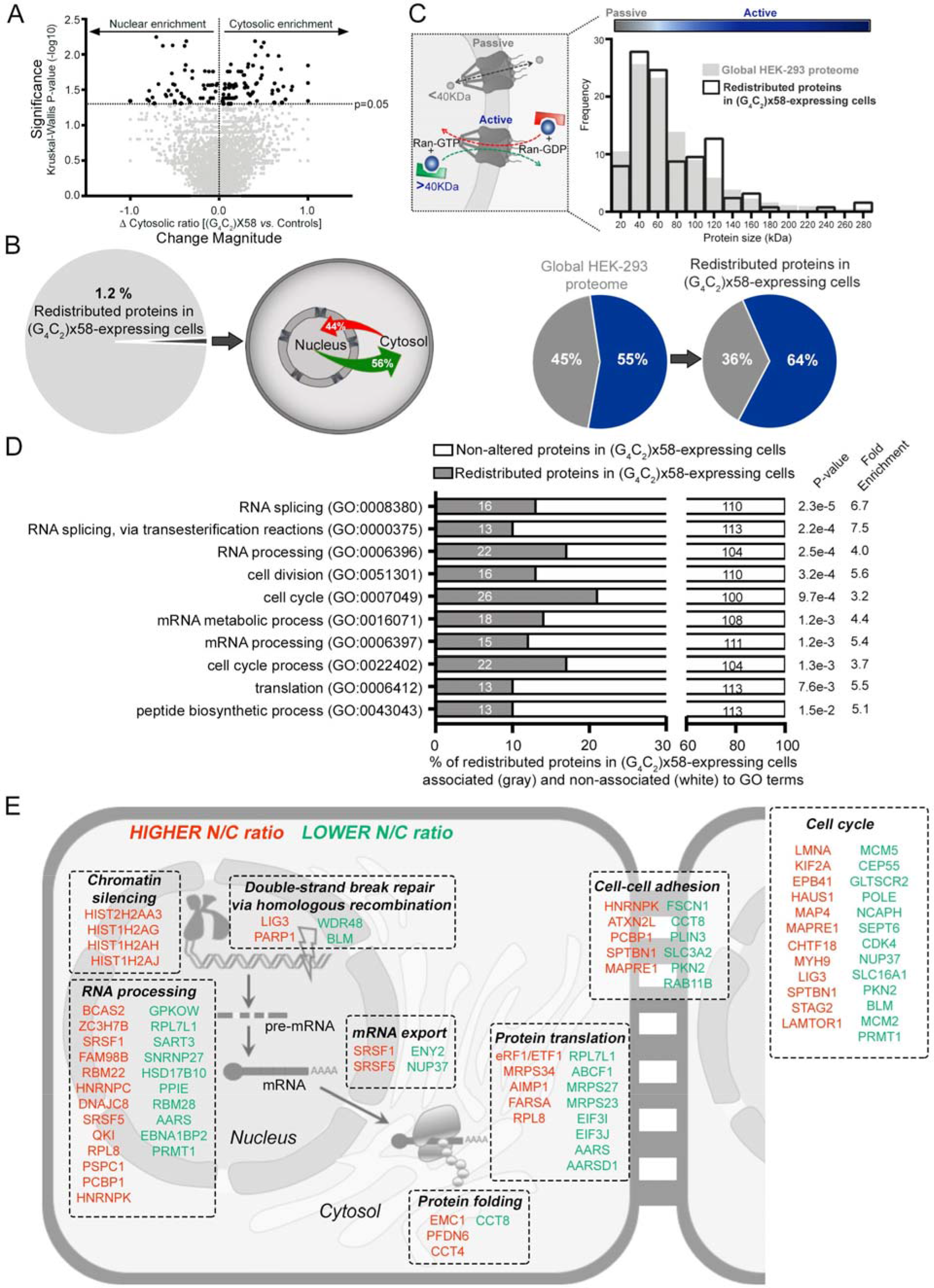
The C9-HRE Causes a Redistribution of Proteins Involved in RNA Metabolism and Protein Translation. (A) Volcano plot representing the magnitude of change of the cytosolic ratio (x-axis) in C9×58-expressing cells compared to both controls (GFP- and C9×8-expressing cells). Y-axis represents the significance of the change (-log10 of the p-value). Proteins that exhibited a significant change were labeled in dark gray. (B) Left: pie chart displaying the percentage of redundant proteins identified by LC-MS/MS that demonstrated statistically significant subcellular redistribution in C9×58-expressing cells compared to controls. Right: Schematic showing the percentage of redistributed proteins that become more nuclear (red) or more cytosolic (green) in C9×58-expressing cells. (C) Top left: graphical representation of size-dependent N/C transport of proteins by passive diffusion or active transport. Top right: histogram showing the size distribution of proteins significantly redistributed in C9×58-expressing cells (black lined bars) compared to the global HEK-293 detected proteome (gray filled bars). Bottom: Pie charts exhibiting the percentage of proteins out of the global HEK-293 cell proteome detected (left) or out of the group of proteins redistributed in C9×58-expressing cells (right), that likely cross the nuclear envelope by passive (gray) or active (blue) transport. (D) Gene ontology (GO) analysis assessed for significant enrichment in biological processes of the 126 unique proteins redistributed in C9×58-expressing cells. Gray bars represent the number of proteins enriched for each pathway relative to the total number of redistributed proteins. P values and fold enrichment values for each cellular process are displayed on the right. (E) Graphical representation of the most significant (p<0.05) GO terms associated with redistributed proteins in C9×58-expressing cells. Proteins with higher (red) or lower (green) N/C ratios are depicted with their corresponding GO term and subcellular localization.

Nucleocytoplasmic exchange of proteins can occur by active transport or passive diffusion through the NPC based on protein mass (Figure 2C, left) (Ma et al., 2012). Interrogating the 126 proteins for size distribution relative to all the proteins identified by MS, revealed a significant enrichment for proteins larger than 40 kDa (p<0.040, 55% to 64%), which likely require active transport through the NPC (Figure 2C, right and bottom). Moreover, while bulk sequence comparisons (Bernhofer et al., 2018) showed no significant enrichment for a NES and/or a NLS (p<0.561, 1.18% to 2.38%), protein-by-protein analysis showed that the majority (79%) of redistributed proteins contained a putative NLS or NES or both (Figure S2B and Table S2). Next, we compared our dataset with previously published proteomic datasets describing distinct transporter cargoes (Kimura et al., 2013a; Kimura et al., 2013b; Kimura et al., 2017; Thakar et al., 2013). We specifically examined cargoes associated with CRM1, which is the major nuclear export receptor, as well as ones allocated to various nuclear import receptors (Importins α/β, -4, -5, -7, -8, -9, -11, and transportins Trn-1, -2, -SR). We found very little overlap between the redistributed proteins and these datasets, likely due to differences in methodology and model systems (Figure S2C and Table S2).

To determine whether our protein targets were translocated as a result of sequestration, we compared them with proteins previously reported to interact with the C9-HRE DPRs and C9-HRE RNA (Boeynaems et al., 2017; Cooper-Knock et al., 2014; Donnelly et al., 2013; Haeusler et al., 2014; Hartmann et al., 2018; Lee et al., 2016; Lin et al., 2016; Lopez-Gonzalez et al., 2016; May et al., 2014; Mori et al., 2013b; Tao et al., 2015; Xu et al., 2013; Yin et al., 2017). We found a strong enrichment of C9-HRE interactors within redistributed proteins relative to all proteins detected (31.29% to 48.41%, p<0.0001) (Figure S2D and Table S2), suggesting sequestration by toxic C9-HRE products contributes in part to the protein redistribution events identified.

We next asked if there was a unifying functional component within the 126 redistributed proteins. Gene ontology (GO) analysis revealed a striking enrichment for proteins involved in RNA metabolism, protein translation and cell cycle dynamics (Figures 2D-E and Table S2). Collectively, our findings indicate that the C9-HRE elicits a significant and highly specific redistribution of proteins that comprise major cellular functions including RNA and protein metabolism.

### Redistributed Proteins are Genetic Modifiers of *C9orf72* Repeat Expansion Toxicity in *Drosophila* Models of Disease

Altered N/C localization of proteins in the context of the C9-HRE could reflect aberrant protein translocation due to sequestration or impaired trafficking, but could also represent a defensive cellular response. To assess the functional significance of redistributed proteins in the context of an intact nervous system, we used transgenic flies expressing 30 or 36 G_4_C_2_ repeats under a GMR-GAL4 driver, which show prominent eye degeneration (Hautbergue et al., 2017; Mizielinska et al., 2014; Xu et al., 2013). We rationally selected six proteins that exhibited a significant magnitude of change in localization (>1.2-fold shift in C9×58-expressing cells relative to both controls), and were representative of the enriched cellular pathways identified in the GO analysis. These included: the mRNA transport protein ENY2, the arginine methyltransferase PRMT1, the RNA-binding protein SRSF1, which was recently linked to *C9orf72* mRNA trafficking (Hautbergue et al., 2017), the nuclear import receptor TNPO3, the chaperone CCT8, and the translation termination factor eRF1 (Figure 3A). We obtained RNAi lines for the fly homologues of these proteins and crossed them with the C9×30 and x36 fly models individually (Figures 3B and S3A). Reducing these proteins in control backgrounds did not elicit eye degeneration (Figure S3B). We found that all six selected proteins were significant genetic modifiers of C9-HRE toxicity *in vivo* (n=12-18 flies/genotype, p<0.01) (Figures 3A-C and S3B-C). Specifically, eRF1, TNPO3, PRMT1 and ENY2 were enhancers in both models as evident from necrotic patches, external eye surface collapse, and depigmentation. CCT8 was an enhancer only in the C9×36 model, and SRSF1 was a strong suppressor of toxicity in both models. This analysis was limited to only six proteins, but there was no indication that the direction of protein redistribution correlated with suppression or enhancement of toxicity in the fly (Figures 3A-C and S3B-C). Collectively, these results suggest that the protein localization analysis we performed highlighted proteins that act as genetic interactors of mutant *C9orf72* toxicity *in vivo*.

**Figure 3.**
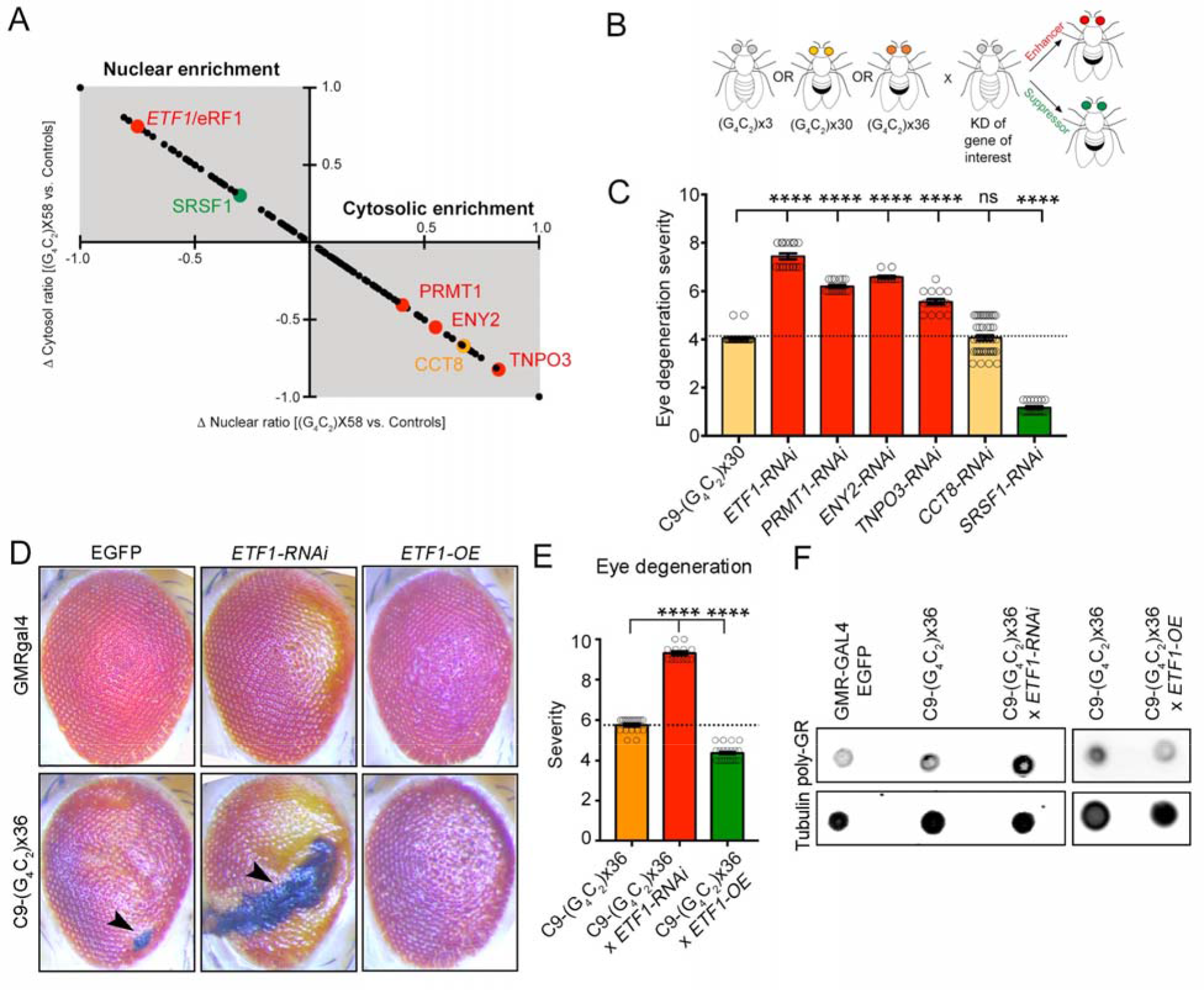
Representative Redistributed Proteins are Genetic Modifiers of C9-HRE Toxicity in *Drosophila* Disease Models. (A) Graph displaying the significantly redistributed proteins, as well as the direction and magnitude of the change observed in C9×58-expressing cells compared to controls. The y-axis represents the change in cytosolic ratio and the x-axis represents the change in nuclear ratio of proteins identified. The upper left gray quadrant includes proteins with a higher N/C ratio, while the lower right quadrant proteins with a lower N/C ratio. Target proteins selected for genetic interaction studies in *Drosophila* are highlighted in green (suppressors), red (enhancers) or orange (no effect) fonts. (B) Schematic illustrating the genetic interaction approach in *Drosophila* models of C9-HRE toxicity. Flies expressing RNAi for selected proteins were crossed with flies expressing 3, 30 or 36 G_4_C_2_ repeats, and the resultant level of toxicity was assessed by the level of suppression (green eye) or enhancement (red eye) of eye degeneration. (C) Bar plots displaying the level of eye degeneration in flies carrying C9×30 repeats with RNAi for *ETF1*, *PRMT1*, *ENY2*, *TNPO3*, *CCT8*, or *SRSF1*. Enhancers and suppressors of toxicity are represented by red and green bars, respectively. The basal level of eye degeneration in C9×30 flies is shown in yellow. Mann-Whitney U test. (D) Representative images of fly eyes from wild-type (GMR-GAL4 x EGFP) or C9-HRE (C9×30 or x36) mutant flies, with endogenous levels (EGFP), knockdown (*ETF1*-RNAi) or overexpression (*ETF1*-OE) of eRF1. Arrow heads indicate the presence of necrotic patches. (E) Bar plots displaying the level of eye degeneration in flies carrying C9-HRE (C9×36 repeats) with endogenous levels, knockdown (*ETF1*-RNAi) or overexpression (*ETF1*-OE) of eRF1. The basal level of eye degeneration in C9×36 flies is shown in orange; suppression and enhancement of C9-HRE toxicity is shown in green and red, respectively. Mann-Whitney U test. (F) Dot blot of poly-GR in control (GMR-GAL4), C9×36, and C9×36 flies with decreased (*ETF1* RNAi) or increased (*ETF1*-OE) eRF1. Tubulin was used as a loading control. In graphs, bars represent mean ± SEM and the dotted lines mark the mean level of eye degeneration in C9-HRE flies.

The eukaryotic translation termination factor 1 (*ETF1*/eRF1), which plays an essential role in RNA metabolism and protein translation (Ivanov et al., 2008; Kashima et al., 2006; Kisselev et al., 2003; Song et al., 2000), was the strongest enhancer of G_4_C_2_ toxicity in both fly models (Figures 3C and S3C). Given the enrichment of these two pathways within the proteins affected by the C9-HRE (Figures 2D-E), we closely examined the effects of eRF1 in C9 *Drosophila* models. Reduction in eRF1 expression enhanced motor dysfunction caused by the HRE compared to controls (n=43-83 flies, p<0.01) (Figure S3D). Conversely, overexpression (Figure S3E) significantly suppressed external eye degeneration, necrotic patches and depigmentation caused by the G_4_C_2_ repeat (n=18-20 flies, p<0.0001) (Figures 3D-E and S3F). Intriguingly, poly-GR levels were relatively higher in the fly eye after knockdown, or lower after overexpression of eRF1 respectively (Figure 3F). These results suggest that eRF1 modulates C9-HRE toxicity *in vivo* by directly or indirectly controlling C9-DPR levels.

### eRF1 is Redistributed in *C9orf72* iPSC-Derived Motor Neurons and Postmortem *C9orf72* ALS Tissue

To validate the shift in eRF1 localization by an alternative approach, we used subcellular fractionation coupled to WB (Figure S4A). We next asked if expressing C9-DPRs alone elicited eRF1 redistribution. Expression of C9×58 (which makes both RNA and DPRs) increased the N/C ratio of eRF1, but expression of codon-optimized constructs (which make each respective DPR but not HRE RNA) did not have a robust effect (Figure S4B-D). These results suggest that eRF1 redistribution is driven by production of G_4_C_2_ repeat RNA and not DPRs.

To determine if our findings on eRF1 redistribution hold significance in a more physiologically relevant disease model system, we differentiated human MNs (Ziller et al., 2018) from iPSCs derived from three individual ALS patients with a *C9orf72* repeat expansion and three healthy controls (Figure S4E-F and Table S3). Subcellular N/C fractionation and WB analysis from differentiated, day 50 MNs, revealed nuclear enrichment of eRF1 in C9 cases (Figure 4A). To validate the redistribution of eRF1 protein at single cell resolution, we used immunocytochemistry (ICC) coupled to confocal microscopy. We found a significantly higher eRF1 N/C ratio in C9-ALS patient MNs relative to controls (day 50, n=3 patients per group, n=3 independent biological replicates; controls *vs* C9: p<0.003, 114% increase ± 34) (Figures 4B-C). Consistent with the role of eRF1 in NMD and translation, the majority of eRF1 protein was cytosolic and in close proximity to the nuclear membrane, but mutant C9 MNs exhibited a higher proportion of nuclear eRF1 signal (Figures 4A-C).

**Figure 4.**
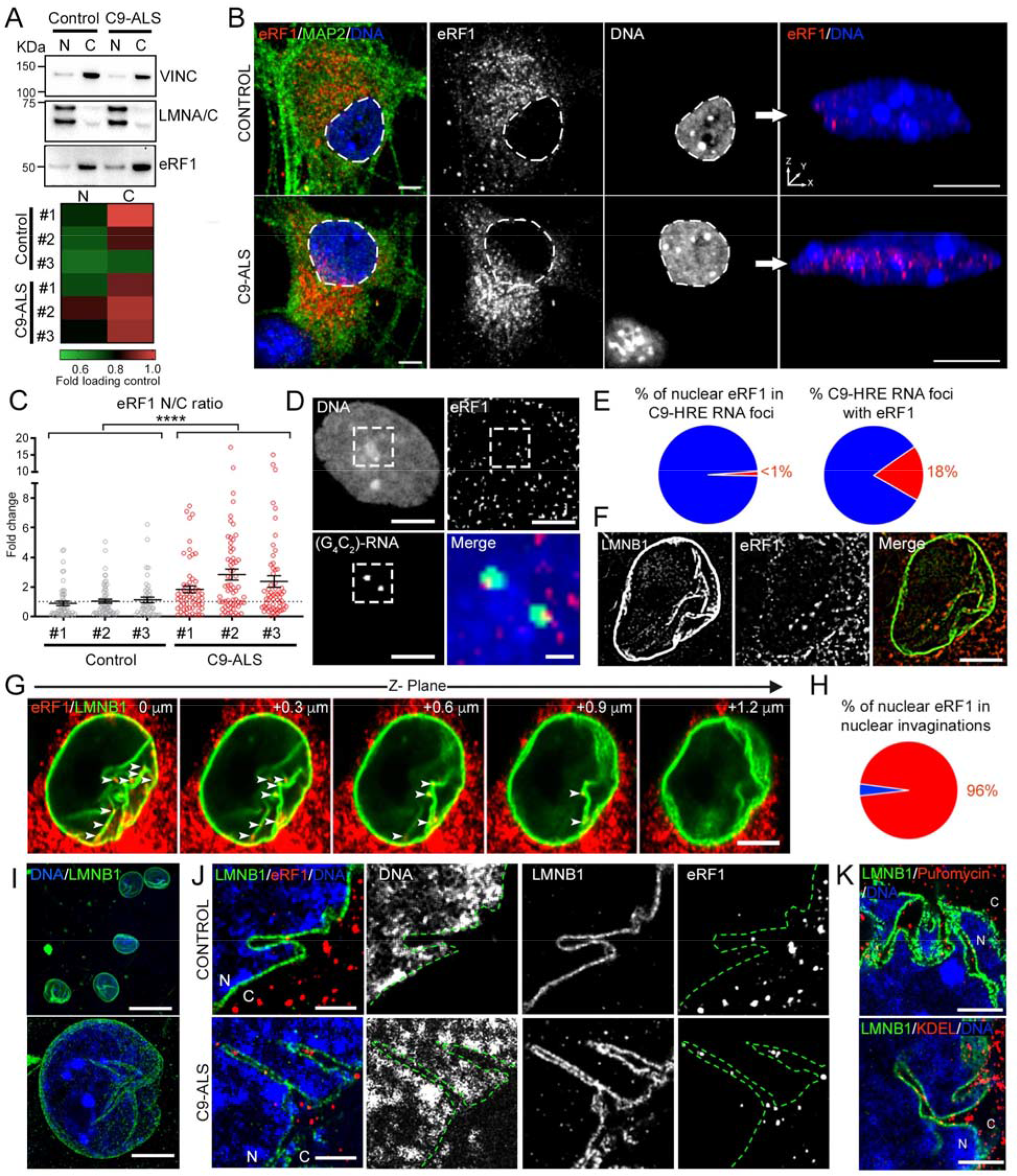
eRF1 is Redistributed in C9-ALS iPSC-Derived MNs. (A) Representative WB bands for eRF1 in nuclear and cytosolic fractions of MN cultures derived from a control and a C9-ALS iPSC line. Bottom: Heat map showing the average eRF1 protein level normalized to subcellular fraction-loading controls (VINCULIN and LMNA/C, for cytosolic and nuclear fractions, respectively) in MNs derived from 3 healthy control and 3 C9-ALS patient iPSC lines (n=5-7 independent differentiations). (B) Left: Representative confocal images of healthy control and C9-ALS patient MNs immunolabeled for eRF1 (red), MAP2 (green) and DNA (blue). Dashed lines mark the nuclear boundary. Right: 3-D reconstruction of control (top) and C9-ALS patient (bottom) MN nuclei shown on the left. (C) Dot plot displaying the fold change in the N/C ratio of eRF1 signal quantified in MNs derived from 3 controls and 3 C9-ALS patient iPSC lines. Individual cells are represented as empty-filled circles, bars represent mean ± SEM, and the dotted line marks the mean N/C ratio in control samples. Mann-Whitney U test. (D) Representative FISH+ICC confocal image of a C9-ALS patient MN nucleus immunolabeled for G_4_C_2_ RNA foci (green), eRF1 (red) and DNA (blue). Dashed lines mark the inset region magnified in the right bottom merged image. (E) Pie charts displaying the percentage of nuclear eRF1 signal that colocalizes with C9-HRE RNA foci (left) and the percentage of C9-HRE RNA foci that colocalize with eRF1 signal (right). (F) Representative structured-illumination microscopy (SIM) image of a C9-ALS patient MN nucleus immunolabeled for eRF1 (red) and LMNB1 (green). (G) Representative serial confocal images of a C9-ALS patient MN nucleus immunolabeled for eRF1 (red) and LMNB1 (green). Arrow heads indicate ERF1+ puncta localized within LMNB1+ nuclear envelope invaginations. (H) Pie chart displaying the percentage of nuclear eRF1 signal that colocalizes with LMNB1+ nuclear envelope invaginations in C9-ALS patient MNs. (I) Regular (top) or expansion (bottom) confocal microscopy images of human iPSC-derived MN nuclei immunolabeled for LMNB1 (green) and DNA (blue). (J) Representative expansion microscopy images of healthy control (top) and C9-ALS patient (bottom) MN nuclei immunolabeled for eRF1 (red), LMNB1 (green), and DNA (blue). (K) Top: Representative expansion microscopy image of a C9-ALS patient MN nucleus immunolabeled for puromycin incorporation (red), LMNB1 (green), and DNA (blue). Bottom: Representative expansion microscopy image of C9-ALS patient MN nucleus immunolabeled for KDEL (red), LMNB1 (green), and DNA (blue). Scale bars: 25 (I), 10 (B left, K), 5 (B right, D, F, G, J), 1 (D inset) μm. N: Nucleus; C: Cytosol.

Using 3D reconstructed confocal images of MN nuclei, we found that eRF1 protein amassed within discrete puncta (Figure 4B, right). We tested whether nuclear accumulation was due to aberrant sequestration by nuclear C9-HRE products (RNA and/or DPRs). Using fluorescence *in situ* hybridization (FISH) coupled to ICC, we found that 18% of nuclear C9-HRE RNA foci colocalized with eRF1 puncta, but less than 1% of nuclear eRF1 puncta colocalized with C9-HRE RNA foci (Figure 4D-E). There was also a negative correlation between nuclear eRF1 and poly-GR/PR signal (Figure S4G). These results suggest C9-HRE RNA/DPRs do not drive the nuclear accumulation of eRF1 by sequestration.

Since neither DPRs nor RNA foci co-localized with nuclear eRF1 puncta, we examined if eRF1 was closely associated with the nuclear membrane, which would result in it being fractionated within the nuclear extract in our MS analysis. Superresolution imaging by structured illumination microscopy (SIM) (Figure 4F) and serial confocal imaging (Figure 4G) revealed a stunning architecture of the nuclear envelope in MNs, with LMNB1+ protrusions within the nucleus, and eRF1 puncta within these structures. Intriguingly, we found that the majority of eRF1 puncta (>96%) colocalized with highly complex LMNB1+ nuclear envelope invaginations (n=3 patients, m=48 neurons per patient) (Figure 4H).

To elucidate whether eRF1 accumulated on the nucleoplasmic or the cytosolic side within nuclear envelope invaginations, we performed expansion microscopy (Asano et al., 2018) coupled to confocal-based imaging (Figure 4I). This approach increased resolution and illustrated that eRF1 resided within the cytosolic side of the nuclear membrane (Figure 4J). These invaginations also demonstrated endoplasmic reticulum (ER) markers, sites of active protein translation (Figure 4K), and almost all had eRF1 protein within them (Figure S4H). Using a knock-in LMNB1-mEGFP reporter iPSC line, we found that all MNs progressively accumulated nuclear envelope invaginations (Figure S4I), suggesting they represent functional structural elements. Critically, C9-ALS patient and control MNs did not show different invagination abundance (Figure S4J-K), but patient MNs contained significantly more eRF1 within these structures (Figure 4B).

To validate these findings in the context of ALS patient neurons *in vivo*, we performed immunohistochemistry (IHC) in postmortem cortical tissue from three *C9orf72* ALS patients and three age-matched controls (Table S3). We found a significantly higher eRF1 N/C ratio in C9-ALS patient MNs (n=3 patients per group, m=28-68 neurons per patient; controls *vs* C9: p<0.0001) (Figures 5A-B and S5). In contrast to patient iPSC-derived MNs, we found a significantly higher abundance of LMNB1+ nuclear envelope invaginations in C9 cases relative to controls (n=3 patients per group, m=31-76 neurons per patient; controls *vs* C9: p<0.0001) (Figure 5C-D). Collectively, these studies show that a proportion of eRF1 becomes redistributed in mutant *C9orf72* patient neurons *in vitro* and *in vivo*.

**Figure 5.**
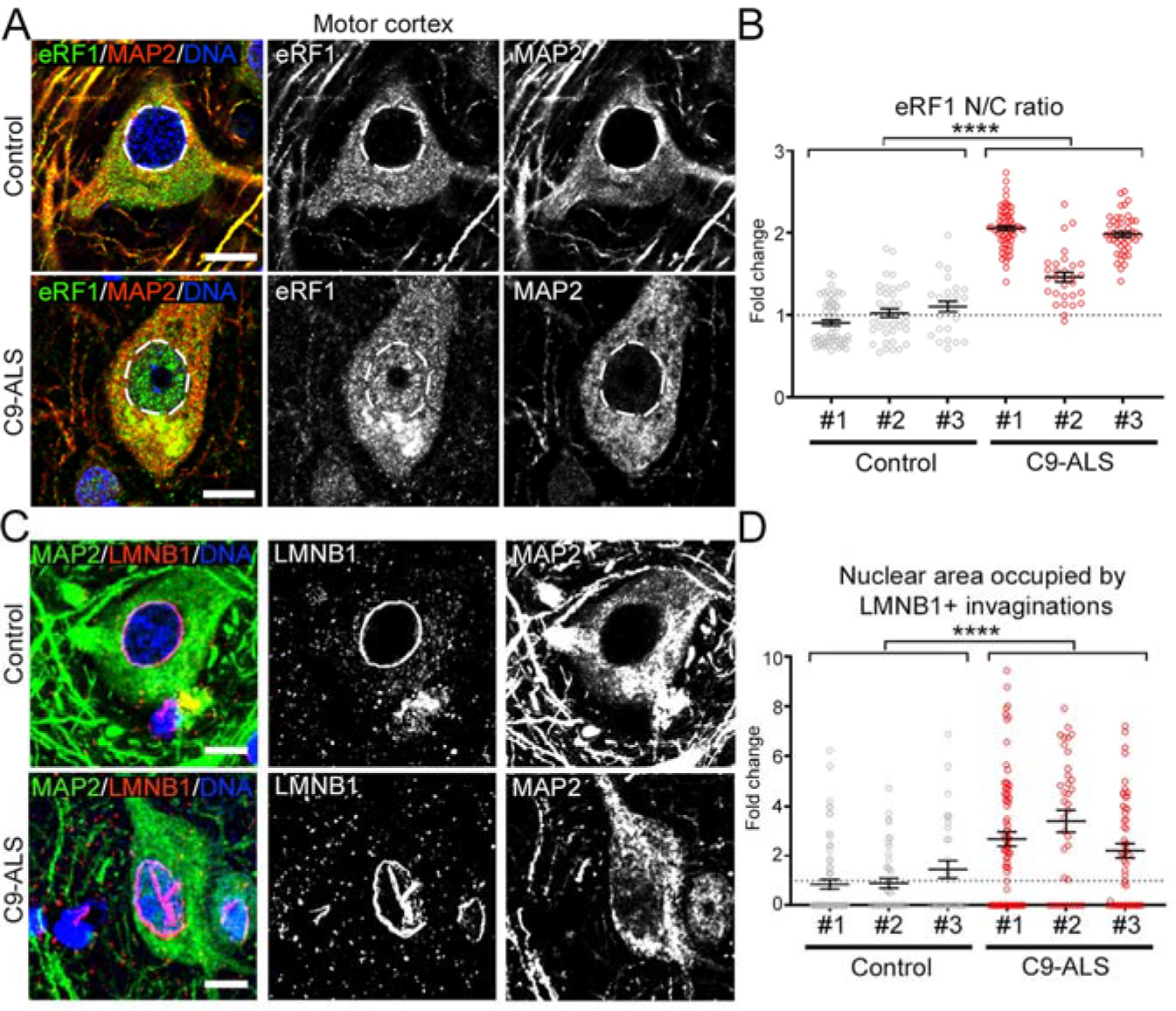
eRF1 is Redistributed in Postmortem C9-ALS Tissue. (A) Representative IHC confocal images of layer V neurons immunolabeled for eRF1 (green), MAP2 (red) and DNA (blue) in motor cortex tissue from non-neurological controls and C9-ALS patients. Dashed lines mark the nuclear boundary. (B) Dot plot displaying the fold change in the N/C ratio of eRF1 signal observed in cortical neurons of 3 non-neurological age-matched controls and 3 C9-ALS patients. Mann-Whitney U test. (C) Representative IHC confocal images of layer V neurons immunolabeled for LMNB1(red), MAP2 (green), and DNA (blue) in motor cortex tissue from non-neurological controls and C9-ALS patients. (D) Dot plot displaying the fold change in the nuclear area occupied by LMNB1+ invaginations in cortical neurons of 3 non-neurological age-matched controls and 3 C9-ALS patients. Mann-Whitney U test. In graphs, bars represent mean ± SEM and the dotted lines mark the mean in control samples. Scale bars: 10 μm.

### eRF1 Mediates a Shift From Protein Translation to mRNA Degradation Through Nonsense Mediated Decay in C9-HRE Expressing Cells

eRF1 plays an essential role in two closely related cellular mechanisms: termination of protein translation and nonsense mediated decay (NMD) of mRNA molecules. During regular termination of protein synthesis, eRF1 recognizes stop codons, binds to the translating ribosome, and triggers cleavage of the nascent peptide (Brown et al., 2015; Kryuchkova et al., 2013; Lind et al., 2017). In cases where eRF1 detects a premature stop or an irregularly spliced mRNA, it triggers its degradation by recruiting the master regulator of NMD, upframeshift protein 1 (UPF1), which becomes phosphorylated (pUPF1), and enlists endonuclease activities (Huang and Wilkinson, 2012; Kashima et al., 2006; Kurosaki et al., 2014; Kurosaki and Maquat, 2016; Wolin and Maquat, 2019).

Given the importance of eRF1 in adjusting the balance of these two cellular pathways, we next asked if its redistribution affected the state of protein translation and NMD. We first used ICC to measure the amount of the methionine analog homopropargylglycine (HPG) incorporated into newly translated proteins at single-cell resolution (Figure S6A). We found a significant reduction in the rate of *de novo* protein synthesis in C9-58-expressing cells relative to controls (n=3 independent biological replicates; GFP *vs*. C9×58: p<0.001, 42% decrease ± 3; and C9×8 *vs*. C9×58: p=0.004, 41% decrease ± 4) (Figure S6B). We next measured the basal rate of protein synthesis in C9 patient and control iPSC-derived MNs (Figure 6A). In accordance with previous reports (Hartmann et al., 2018; Kanekura et al., 2016; Kwon et al., 2014; Lee et al., 2016), C9-ALS patient MNs exhibited significantly reduced protein synthesis relative to controls (day 50, n=3 patients *vs* 3 control, n=6-10 independent biological replicates; p<0.01, 18% decrease ± 3) (Figure 6B). Critically, eRF1 knockdown led to a significant reduction but not complete ablation of translation in healthy MNs (siScr *vs* si*ETF1* in n=1 control, n=3 independent biological replicates, p<0.0001, 60% decrease ± 5), confirming that eRF1 is responsible for translation under control conditions (Figures 6C and S6C). However, eRF1 knockdown had a more moderate effect in C9 MNs, likely because translation is already significantly impaired (siScr *vs* si*ETF1* in n=2 patients, n=3 independent biological replicates, p=0.336, 22% decrease ± 4) (Figure 6C). Since activation of the integrated stress response (ISR) can inhibit protein synthesis (Harding et al., 2003) and trigger DPR translation (Westergard et al., 2019), we addressed whether it was contributing to this translational defect. We did not find any strong evidence for ISR activation in patient MNs based on the level of three ISR markers (Figure S6D). Additionally, treatment with ISRIB, which reverses the effects of eIF2α phosphorylation, did not restore protein translation levels in C9 MNs (Figure S6E).

**Figure 6.**
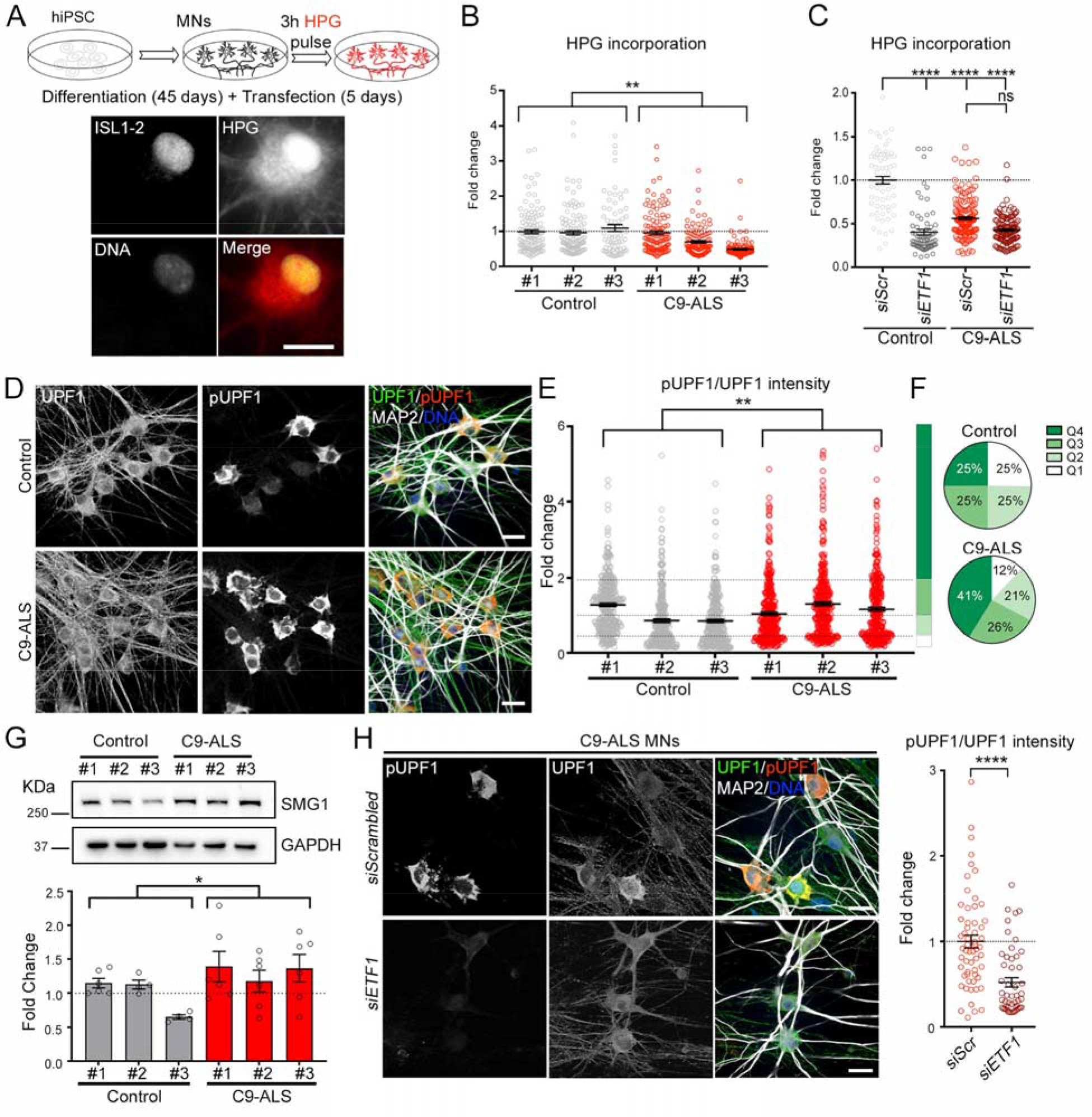
Protein Translation and mRNA Degradation Through NMD in C9-HRE Expressing Cells. (A) Top: schematic of *de novo* protein translation assay performed in iPSC-derived MNs. Bottom: representative image of HPG fluorescent labeling of *de novo* protein synthesis in an ISL1-2+ MN. (B) Dot plot displaying the level of HPG incorporation in MNs derived from 3 control and 3 C9-ALS iPSC lines. Kruskal-Wallis test. (C) Dot plot displaying the level of HPG incorporation in MNs transfected with scrambled (siScr) or *ETF1* siRNAs. MNs derived from 1 control and 2 C9-ALS iPSC lines. Kruskal-Wallis test; ns=not significant. (D) Representative confocal images of healthy control and C9-ALS patient MNs immunolabeled for total UPF1 (green), pUPF1 (red), and DNA (blue). (E) Dot plot displaying the fold change in pUPF1/UPF1 signal in control and C9-ALS MNs. (Mann-Whitney U test. (F) Pie charts displaying NMD activation levels defined by pUPF1/UPF1 quartiles (Q1-4) observed in control (top) and C9 (bottom) MNs. (G) Top: Representative WB for SMG1 in MN cultures derived from 3 control and 3 C9-ALS iPSC lines. GAPDH was used as a loading control. Bottom: bar plots showing the fold change in the SMG1/GAPDH ratio in control and C9-ALS MN cultures. T test. (H) Left: Representative confocal images of C9-ALS patient MNs transfected with scrambled or *ETF1* siRNAs and immunolabeled for total UPF1 (green), pUPF1 (red), and DNA (blue). Right: dot plot displaying the fold change in pUPF1/UPF1 signal in C9-ALS MNs transfected with scrambled or *ETF1* siRNAs. n=3 independent differentiations; Mann-Whitney U test. In graphs, empty-filled circles represent individual cells (B, C, E, H) or biological replicates (G). Bars represent the mean ± SEM and dotted lines mark the mean level in control samples (B, C, G, H) or Q1-4 quartile limits (E). Scale bars: 25 (D, H), 20 (A) μm.

In contrast to reduced protein synthesis, C9-ALS patient MNs exhibited an induction of NMD as measured by the levels of pUPF1 protein (day 50, n=3 patients per group, n=4 independent biological replicates; controls *vs* C9: p<0.007, 38% increase ± 13) (Figure 6D-F). Critically, C9-ALS cultures showed a striking increase in the proportion of MNs with high pUPF1 levels relative to controls, indicating hyperactivated NMD (from 25% in controls to 41% in patients, n>217 MNs per individual) (Figure 6F). We also examined the level of SMG1, which is the canonical kinase that phosphorylates UPF1, and found a significant increase in C9-ALS MN cultures (n=6 independent biological replicates; controls *vs* C9: p<0.05, 31% increase ± 11) (Figure 6G). Importantly, NMD activation was eRF1-dependent as its knockdown in HEK-293 cells (Figure S6F) or patient-derived MNs (Figure 6H), significantly diminished pUPF1 levels (p<0.0001). We then used a reporter system (Nickless et al., 2014) that monitors NMD efficiency based on the ratio of a transcript with a premature termination codon (PTC) to one without (Figure S6G-I). We found that C9-58 expression triggered NMD as the ratio of PTC to control transcript showed a small but significant decrease relative to C9-8-expressing cells (n=3 independent biological replicates; C9×8 *vs*. C9×58: p=0.044) (Figure S6J). Critically, the degradation of the PTC transcript was eRF1-dependent, as this ratio significantly increased after eRF1 knockdown (n=2 independent biological replicates; C9×58: siScr *vs*. siETF1 p<0.0001) (Figure S6K-L).

### *C9orf72* mRNA is Targeted for Degradation by Nonsense Mediated Decay

Collectively, our data suggest that the C9-HRE elicits an eRF1-mediated shift from protein translation to NMD-dependent mRNA degradation (Figure 6). This shift likely confers a protective effect as eRF1 reduction enhances poly-GR production and C9-HRE toxicity, while eRF1 overexpression suppresses C9-HRE toxicity and poly-GR production in *Drosophila* C9 models (Figure 3). The C9-HRE also triggers eRF1 accumulation in nuclear envelope invaginations (Figure 4). Importantly, we found evidence for protein synthesis within these invaginations based on detection of puromycin, ER markers, as well as colocalization of eRF1 and pUPF1 (Figures 4K and S7A-B), suggesting that a proportion of mRNA surveillance coupled to protein translation occurs within these structures.

It is likely that the reported C9-HRE-dependent retention of the first intron of *C9orf72* in combination with the high GC content of the HRE, earmarks the expanded transcript as a target for NMD-dependent degradation (Sznajder et al., 2018; Colombo et al., 2017; Imamachi et al., 2017; Zhang et al., 2009). To test this hypothesis, we employed several approaches. Given that NMD degrades mRNAs as they exit the nuclear pore complex (Kurosaki and Maquat, 2016), we analyzed the association between eRF1 localization and C9-HRE RNA foci, a prototypical pathological marker of *C9orf72*-associated ALS. We found that C9-ALS patient MNs with RNA foci had a significantly higher eRF1 N/C ratio than MNs without foci (Figure 7A), suggesting that accumulation of eRF1 within invaginations may degrade or inhibit export of the expanded transcript. We next asked whether the absence of eRF1 would affect the number or localization of RNA foci. We found that eRF1 knockdown led to a dramatic reduction of nuclear foci and a concomitant increase in cytosolic foci in C9-ALS MNs (day 50, n=3 patients, n=3 independent biological replicates; siScr *vs*. siETF1 nuclear foci: p<0.0001; cytosolic foci: p<0.0001) (Figure 7B-C). eRF1 knockdown, which abolishes NMD surveillance, also resulted in a lower N/C ratio for all poly(A) RNA, to levels equal to the ones seen in control MNs (Figure 7D-E). These data suggest that the reported nuclear retention of RNA in C9-ALS MNs (Freibaum et al., 2015), is partly mediated by an eRF1-dependent induction of the NMD pathway.

**Figure 7.**
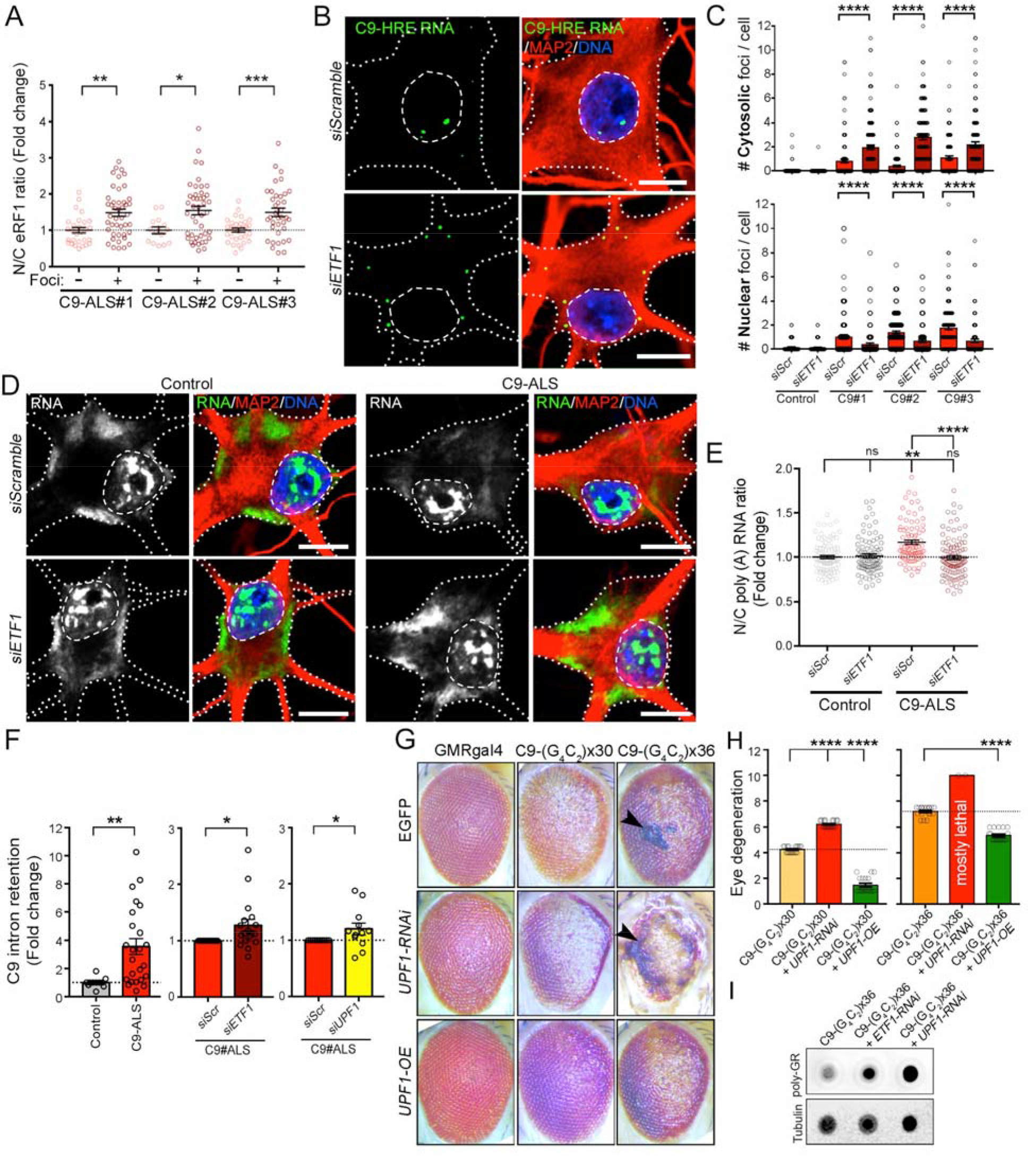
C9orf72 mRNA is Targeted for Degradation by NMD. (A) Dot plot displaying the fold change in the N/C ratio of eRF1 signal quantified in C9-ALS MNs with (+) or without (-) C9-HRE RNA foci. n=2 independent differentiations; Mann-Whitney U test; p*<0.05; **<0.01; ***<0.001. (B) Representative FISH+ICC confocal images of C9-ALS patient MNs transfected with scrambled or *ETF1* siRNA and immunolabeled for G_4_C_2_ RNA foci (green), MAP2 (red) and DNA (blue). Dashed lines mark the nuclear boundary and dotted lines mark the neuronal soma. (C) Bar plots showing the number of nuclear (bottom) and cytosolic (top) C9-HRE RNA foci per cell in MNs derived from 1 control and 3 C9-ALS iPSC lines, transfected with scrambled or *ETF1* siRNAs. Wilcoxon signed rank. (D) Representative FISH+ICC confocal images of healthy control and C9-ALS patient MNs transfected with scrambled or *ETF1* siRNAs, and immunolabeled for poly(A) RNA (green), MAP2 (red) and DNA (blue). Dashed lines mark the nuclear boundary and dotted lines mark the neuronal soma. (E) Dot plot showing the N/C ratio of poly(A) RNA in control and C9-ALS MNs transfected with scrambled or *ETF1* siRNAs. n=2 independent differentiations; Kruskal-Wallis test, p=ns: not significant; **<0.01, ****<0.0001;. (F) Left graph: Bar plots showing fold change in the level of *C9orf72*-intron retention in MN cultures derived from 3 control and 3 C9-ALS iPSC lines as measured by semi-quantitative RT-PCR. Bar plots comparing intron retention in C9-ALS MNs transfected with scrambled (Scr) *vs. ETF1* (middle graph) or *vs. UPF1* (right graph) siRNAs. Mann-Whitney (left and right graphs), and t test (middle graph). (G) Representative images of fly eyes from wild-type (GMR-GAL4 x EGFP) or C9-HRE (C9×30 or x36) mutant flies, with endogenous levels (EGFP), knockdown (*UPF1*-RNAi) or overexpression (*UPF1*-OE) of UPF1. Arrow heads indicate the presence of necrotic patches. (H) Bar plots displaying the level of eye degeneration in flies carrying C9-HRE (C9×30 or x36 repeats) with endogenous levels, knockdown (*UPF1*-RNAi) or overexpression (*UPF1*-OE) of UPF1. The basal level of eye degeneration in C9×30 or x36 flies is shown in yellow and orange respectively; suppression and enhancement of C9-HRE toxicity is shown in green and red, respectively. Mann-Whitney U test. (I) Dot blot of poly-GR in C9×36, and C9×36 flies with decreased eRF1 (*ETF1*-RNAi) or UPF1 (*UPF1*-RNAi). Tubulin was used as a loading control. In graphs, empty-filled circles represent individual cells (A, C, E), biological replicates (F) or analyzed flies (H). Bars represent the mean ± SEM, and dotted lines mark the mean levels in control/reference samples. Scale bars: 10 μm.

To directly address the effects of eRF1 on the stability of G_4_C_2_ repeat RNA (including the soluble fraction that may not end up in discreet foci), we measured the ratio of mutant to wild type C9 transcript based on intron retention (Figure S7C). We first established that C9-ALS patient MNs exhibited significantly higher intron retention than control MNs (day 50, n=3 patients per group, n=1-8 independent biological replicates; controls *vs* C9: p=0.0046) (Figures 7F and S7D, F), which was exacerbated upon eRF1 knockdown (n=3 patients, n=6 independent biological replicates; siScr *vs*. si*ETF1*: p=0.014) (Figures 7F and S7D-F). These results accord with the higher levels of poly-GR upon eRF1 knockdown found *in vivo* (Figure 3F), as higher cytosolic C9-HRE levels may allow for more DPR translation. Critically, UPF1 reduction, which abolishes NMD also exacerbated intron retention, suggesting that the expanded transcript may be an NMD target (n=3 patients, n=3 independent biological replicates; siScr *vs*. siUPF1: p=0.011) (Figures 7F and S7D-F).

Finally, to directly assess the significance of NMD and UPF1 in mutant *C9orf72* toxicity *in vivo,* we crossed UPF1 overexpressing and RNAi lines with C9-HRE fly models. In control flies these crosses had minimal effects on eye degeneration (Figures 7G and S7G). UPF1 reduction significantly enhanced eye degeneration in the C9×30 fly model (n=18 flies/genotype, p<0.01), and was overly toxic in the C9×36 fly model, with the majority of flies dying at the pupal stage (Figures 7G-H and S7H). Conversely, UPF1 overexpression significantly ameliorated eye degeneration in both C9-HRE fly models (n=18 flies/genotype, p<0.0001) (Figures 7G-H). In accordance with these effects and a putative UPF1 role in targeting the expanded C9 RNA for NMD degradation, we found that UPF1 reduction led to higher poly-GR levels *in vivo*, similar to results upon *ETF1* knockdown (Figure 7I). In summary, these results indicate that eRF1 mediates an UPF1-dependent degradation of the expanded *C9orf72* transcript.

## DISCUSSION

The transport of mRNA and proteins between the nucleus and cytoplasm has emerged as a critical pathway in neurodegenerative diseases (Eftekharzadeh et al., 2018; Gasset-Rosa et al., 2017; Grima et al., 2017; Woerner et al., 2016; Zhang et al., 2016). We developed an approach to assess how the C9-HRE affects the subcellular N/C distribution of proteins. We found a small but highly specific redistribution of proteins in C9×58-expressing cells, the majority of which were associated with RNA metabolism and translation. While the proteomic analysis of the N/C distribution of proteins provided new insight, the cellular platform we used was selected to interrogate acute effects of the mutation. It is possible that steady state protein localization changes in neurons chronically exposed to the expansion may be different. Experiments with distinct repeat expansion mutations revealed that they all caused a global shift towards a more cytosolic proteome, yet the identities of the redistributed proteins were different in each case. Future examination of the N/C proteome in distinct neuronal subtypes may elucidate neuron-specific protein localization effects that repeat expansion mutations associated with different neurodegenerative diseases may have.

Our findings strengthen the hypothesis that a major component of mutant *C9orf72* toxicity is related to mRNA processing defects. Although the repeat expansion in *C9orf72* is by far the largest genetic cause of ALS and FTD (DeJesus-Hernandez et al., 2011; Renton et al., 2011), the extent to which C9 pathophysiological mechanisms are implicated in other ALS cases remains unclear. ALS genetics highlight RNA metabolism as a principle disease mechanism with causal mutations identified in a number of RNA-binding proteins such as TDP-43, FUS, MATR3 and HNRNPA1 (Taylor et al., 2016). Moreover, a number of studies have extensively characterized RNA metabolism defects in ALS (Weskamp and Barmada, 2018). These include RNA instability (Tank et al., 2018) and defective RNA splicing in mutant TDP-43 and FUS models (Lagier-Tourenne et al., 2012; Polymenidou et al., 2011; Reber et al., 2016), as well as within postmortem patient brain samples from *C9orf72* and sporadic ALS cases (Prudencio et al., 2015).

We found that C9-ALS patient MNs exhibit NMD hyperactivity in accordance with a recent study, which showed that ALS-causing *FUS* mutations also lead to NMD hyperactivity (Kamelgarn et al., 2018). The activation may be triggered by the expression of the expanded *C9orf72* transcript directly or by broader mRNA processing defects that occur downstream of the mutation (Prudencio et al., 2015). Specifically, sequestration of mRNA metabolism proteins within C9-HRE foci and C9-HRE DPR aggregates (Boeynaems et al., 2017; Hautbergue et al., 2017; Kwon et al., 2014; Lee et al., 2016; Mori et al., 2013b; Xu et al., 2013; Yin et al., 2017), may result in extensive splicing errors that trigger mRNA surveillance pathways including NMD (Zhang et al., 2009).

Our work highlights eRF1 and UPF1 as strong modulators of *C9orf72*-associated pathophysiology and provides a functional link between the C9-HRE and the NMD pathway. UPF1 has previously been implicated in ALS disease mechanisms related to FUS and TDP-43 toxicity (Armakola et al., 2012; Barmada et al., 2015; Jackson et al., 2015; Ju et al., 2011). It was reported to mediate the suppressive effects of DBR1 on TDP-43 toxicity in yeast (Armakola et al., 2012), while its overexpression was beneficial for mutant TDP-43 and FUS rodent neurons *in vitro* (Barmada et al., 2015; Ju et al., 2011), as well as in a rat model of TDP-43 toxicity *in vivo* (Jackson et al., 2015). More recently, Xu et al., reported that poly-GR/PR inhibit NMD, and in accordance with our findings, that UPF1 overexpression or pharmacological NMD activation suppressed DPR toxicity (Xu et al., 2019).

The canonical site of NMD has been a subject of debate (Kurosaki and Maquat, 2016), with most models suggesting that it occurs in the cytoplasm near the nuclear membrane and immediately upon mRNA export (Belgrader et al., 1994; Trcek et al., 2013). Our findings suggest that, at least in MNs, NMD can occur within nuclear envelope invaginations, which contain ER proteins, eRF1, pUPF1, and are sites of active translation. Tubular invaginations of the nuclear envelope have been reported in a wide range of cell types, and are prominent in dividing cells and cancer tissue (Malhas et al., 2011). While the mechanistic relevance of these protrusions remains unclear, they have been associated with DNA repair (Legartova et al., 2014) and mRNA transport (Schoen et al., 2017). These invaginations are not exclusive to C9-ALS patient MNs, nor do they represent an *in vitro* artifact. We propose that they serve to extend cytosolic function related to RNA metabolism and translation in closer proximity to the nucleus, perhaps to increase the efficiency of these critical cellular processes by shortening the distance between the nucleus and cytosol, or by increasing the N/C interaction area. This functional role would be particularly relevant to postmitotic neurons, which may heavily rely on the ability to rapidly and dynamically regulate RNA metabolism and compartmentalized protein translation. Future investigations will need to address what these invaginations contain, their functional relevance, and if they truly represent a beneficial neuronal response or a pathological consequence of aging and disease.

The accumulation of eRF1 and pUPF1 within nuclear envelope invaginations suggest a model where NMD safeguards neurons from toxic RNA molecules, such as the expanded *C9orf72,* by inhibiting transcript export from the nucleus and the ensuing cytosolic translation into toxic C9-DPRs (Figure S8), through an as yet undefined mechanism. The potential therapeutic relevance of this model could be explored either through small molecules that would activate NMD, or directly through gene therapy approaches. The cytosolic accumulation of TDP-43 as well as erroneous mRNA metabolism is a unifying pathological feature amongst ALS patients (Tank et al., 2018; Taylor et al., 2016), suggesting that the NMD pathway and its regulatory elements may represent a therapeutic approach that could be broadly relevant for ALS and FTD.

## ACKNOWLEDGEMENTS

We are grateful to the following funding sources: US National Institutes of Health (NIH) / National Institute on Neurological Disorders and Stroke (NINDS) and National Institute on Aging (NIA) R01NS104219 (EK), NIH/NINDS grants R21NS111248 (EK), R21NS107761 (EK/JNS), Les Turner ALS Foundation (EK), Muscular Dystrophy Association (EK and UBP), ALS Association (UBP), Robert Packard Center for ALS at Johns Hopkins (UBP), NIH R01NS081303 (UBP), R21 NS098379, NS101661, NS100055 (UBP) and NIH/NINDS R01AG061787 (JNS). We thank Alfred W Rademaker for advice on statistical analysis. JAO is a postdoctoral fellow of The French Muscular Dystrophy Association (AFM-Téléthon). EK is a member of the Simpson Querrey Institute for BioNanotechnology, a Les Turner ALS Research Center Investigator and a New York Stem Cell Foundation – Robertson Investigator.

## AUTHOR CONTRIBUTIONS

Conceptualization, JAO, ELD and EK; Methodology and Validation, JAO, ELD, SK, JNS, UBP and EK; Formal Analysis, JAO, ELD and EK; Investigation, JAO, ELD, SK, MS, LT, HSS, EAH, TYE, YT, TFG, CJD, TS, JNS, UBP and EK; Resources, TYE, TFG, CJD, TS, JNS, UBP and EK; Writing – Original Draft, JAO, ELD and EK; Writing – Review & Editing, JAO, ELD, SK, MS, LT, HSS, EAH, TYE, YT, TFG, CJD, TS, JNS, UBP and EK; Supervision and Project administration, EK; Funding Acquisition, JNS, UBP and EK.

## STAR METHODS

### LEAD CONTACT AND MATERIALS AVAILABILITY

Further information and requests should be directed to and will be fulfilled by the Lead Contact, Evangelos Kiskinis.

### EXPERIMENTAL MODEL AND SUBJECT DETAILS

#### Cell culture conditions

HEK-293FT cells were grown in DMEM (Corning) supplemented with Glutamax (Gibco) and 10% fetal bovine serum (FBS, Gibco). HEK-293FT cells were dissociated by incubating for 5 min with Trypsin-EDTA (Gibco) at 37°C. Stem cells were maintained on Matrigel (BD Biosciences) with mTeSR1 media (Stem Cell Technologies) and passaged on a weekly basis using 1mM EDTA or Accutase (Sigma). All cell cultures were maintained at 37°C, 5% CO_2_ without antibiotics and tested on a monthly basis for mycoplasma. All cell lines utilized in the study were mycoplasma-free.

#### Stem cell cultures and motor neuron differentiation

iPSCs were dissociated with Accutase and plated with mTESR1 and 10µM ROCK inhibitor (Y-27632, DNSK International) at a density of 100,000 cells/cm^2^. The next day, media was replaced with N2B27 medium (50% DMEM:F12, 50% Neurobasal, supplemented with NEAA, Glutamax, N2 and B27; Gibco) supplemented with a cocktail of small molecules that induces the generation of spinal neural progenitors: 10µM SB431542 (DNSK International), 100nM LDN-193189 (DNSK International), 1µM Retinoic Acid (RA, Sigma) and 1µM of Smoothened-Agonist (SAG, DNSK International). Culture medium was changed daily for 6 days and then was switched to N2B27 medium supplemented with 1µM RA, 1µM SAG, 5µM DAPT (DNSK International) and 4µM SU5402 (DNSK International) to generate postmitotic spinal MNs. Cultures were fed daily with this media for seven days and then dissociated using TrypLE Express (Gibco) supplemented with DNase I (Worthington). Neurons were plated onto culture dishes precoated with Matrigel (BD Biosciences) and fed every 2 days with Neurobasal medium supplemented with NEAA, Glutamax, N2, B27, Ascorbic acid (0.2µg/ml) and BDNF, CNTF and GDNF (10ng/mL, R&D systems). For imaging analysis, MNs were seeded on preplated mouse glia monolayers.

### METHOD DETAILS

#### Overexpression and knockdown experiments *in vitro*

For overexpression experiments, 40% confluent HEK-293 cells were transfected with HilyMax transfection reagent (Dojindo Molecular Technologies) according to manufacturer guidelines. Briefly, DNA was mixed with HilyMax (1µg DNA:3µL HilyMax ratio) in Opti-MEM medium (Gibco) and incubated for 15 min at room temperature (RT) before being added to cells. Cells were incubated with transfection mixture for 4hrs at 37°C and then media was replaced. Analyses made on transfected cells were performed 48 to 72hrs after transfection. For transient knockdown experiments, pre-designed siRNAs (Ambion Silencer Select) were transfected using Lipofectamine RNAiMAX (Invitrogen) according to manufacturer guidelines. Briefly, for 2 wells of a 24-well plate with 0.5-2 x 10^5^ cells/well, 10pmol of siRNA was mixed with 3µL of Lipofectamine RNAiMAX reagent in 100µL Opti-MEM medium and incubated for 15 min at RT. 400µL of cell culture medium was added and 250µL was distributed per well. Analyses of siRNA transfection experiments were performed 3-4 days after transfection. For lentiviral transductions, iPSC-derived MNs were infected with a previously tittered viral burden for 48hrs, changing the media and adding fresh virus 24hrs after the initial infection. Transduced cells were analyzed 10-14 days after infection.

#### Plasmids

For Mass Spectrometry experiments, HEK-293 cells were transfected with the following DNA constructs: pcDNA3.1 plasmids containing GFP or (G_4_C_2_)x58-GFP provided by J. Paul Taylor’s lab; pEGFP plasmids containing HTT exon1 with (CAA CAGCAGCAACAGCAA)n repeats amounting to either 25Q or 72Q repeats provided by Dimitri Krainc’s lab; pLEX plasmids containing Atx1-32Q or Atx1-84Q provided by Puneet Opal’s lab. The pcDNA3.1 plasmids containing GFP, (GA)x50-, (GR)x50-, (PA)x50-, or (PR)x50-GFP, in which alternative codons were used to generate dipeptide repeat proteins without generating the (GGGGCC)n transcript were originally made by Davide Trotti (Wen et al., 2014) and modified by Christopher Donnelly. We also used this plasmid to subclone the GFP and (GR)x50-GFP sequences into a lentiviral vector (CD510B) for MN transductions. The pLenti-c-Myc-DDK plasmid containing NLS-tdTomato-NES was provided by Jeffrey Rothstein (Zhang et al., 2015). The pBS-TCR(PTC)-TCR(WT) NMD reporter plasmid was provided by Zhongsheng You (Nickless et al., 2014).

#### Flow cytometry and nucleocytoplasmic fractionation

Cell cultures were dissociated with trypsin and resuspended in media containing DMEM without phenol red (Gibco) and 2% FBS before flow cytometry cell sorting on a BD FacsAria SORP 6-Laser instrument. Cells were collected in PBS supplemented with protease inhibitors (Millipore). Cell suspensions were immediately spun down and subcellular fractionation was performed using NE-PER nuclear and cytoplasmic extraction reagents (Thermo Scientific). Briefly, cells were harvested using a hypertonic detergent solution that disrupted plasma membranes, leaving nuclear membranes intact. This lysate was spun down and supernatant was collected as the cytosolic protein fraction. Nuclear proteins were isolated from the remaining pellet using high salt extraction buffer. The obtained nuclear and cytosolic fractions were used for Western blot and mass spectrometry analysis.

#### Liquid Chromatography Mass Spectrometry (LC-MS/MS)

Nuclear and cytosolic extracts were subjected to methanol and chloroform precipitation. The precipitated protein pellets were solubilized first in 100μl of 8M urea for 30min, and next in 100μl of 0.2% ProteaseMAX (Promega) for 2hrs. The protein extracts were reduced and alkylated as described previously (Chen et al., 2008), followed by the addition of 300μl of 50mM ammonium bicarbonate, 5μl of 1% ProteaseMAX, and 20μg of sequence-grade trypsin (Promega). Samples were digested overnight in a 37°C thermomixer (Eppendorf). We used 3μg of peptides in each Orbitrap Fusion MS analysis.

For Orbitrap Fusion Tribrid MS analysis, the tryptic peptides were purified with Pierce C18 spin columns and fractionated with two ACN concentrations (30% and 70%). Three micrograms of each fraction were auto-sampler loaded with a Thermo Fisher EASY nLC 1000 UPLC pump onto a vented Acclaim Pepmap 100, 75μm x 2 cm, nanoViper trap column coupled to a nanoViper analytical column (Thermo Fisher m, C18, 0.075mm, 500mm) with stainless steel emitter tip 164570, 3μ, 100AD assembled on the Nanospray Flex Ion Source with a spray voltage of 2000 V. Buffer A contained 94.785% H_2_O with 5% ACN and 0.125% FA, and buffer B contained 99.875% ACN with 0.125% FA. The chromatographic run was for 4hrs in total with the following profile: 0–7% for 7min, 10% for 6min, 25% for 160min, 33% for 40min, 50% for 7, 95% for 5min, and 95% again for 15min, respectively. Additional MS parameters include: ion transfer tube temp 300°C, easy-IC internal mass calibration, default charge state = 2 and cycle time = 3s. Detector type set to Orbitrap, with 60K resolution, with wide quad isolation, mass range normal, scan range = 300–1500m/*z*, max injection time = 50ms, AGC target = 200,000, microscans = 1, S-lens RF level = 60, without source fragmentation, and datatype = positive and centroid. MIPS was set as on, included charge states = 2–6 (reject unassigned). Dynamic exclusion enabled with *n* =1 for 30 and 45s exclusion duration at 10ppm for high and low. Precursor selection decision = most intense, top 20, isolation window = 1.6, scan range = auto normal, first mass = 110, collision energy 30%, CID, Detector type = ion trap, Orbitrap resolution = 30K, IT scan rate = rapid, max injection time = 75ms, AGC target = 10,000, Q = 0.25, inject ions for all available parallelizable time.

#### Tandem mass spectra analysis

Samples of 4 independent biological replicates, which included both nuclear and cytosolic fractions of each of the 3 transfected conditions (GFP, 8 and 58 x G_4_C_2_), were analyzed independently by LC-MS/MS. The spectral files from all replicates were pooled for a single database search. Spectrum raw files were extracted into MS1 and MS2 files using in-house program RawXtractor or RawConverter (http://fields.scripps.edu/downloads.php) (He et al., 2015). The tandem mass spectra were searched against UniProt human protein database (downloaded on 12-19-2015; UniProt Consortium, 2015) and matched to sequences using the ProLuCID/SEQUEST algorithm (ProLuCID version 3.1;(Eng et al., 1994; Xu et al., 2006)) with 50ppm peptide mass tolerance for precursor ions and 600ppm for fragment ions. The search space included all fully and half-tryptic peptide candidates that fell within the mass tolerance window with no miscleavage constraint, assembled, and filtered with DTASelect2 (version 2.1.3) (Cociorva et al., 2007; Tabb et al., 2002) through Integrated Proteomics Pipeline IP2 version 3, Integrated Proteomics Applications (http://www.integratedproteomics.com.). In order to estimate peptide probabilities and false-discovery rates (FDR) accurately, we used a target/decoy database containing the reversed sequences of all the proteins appended to the target database (Peng et al., 2003). Each protein identified was required to have a minimum of one peptide of minimal length of six amino acid residues; however, this peptide had to be an excellent match with an FDR = 0.001 and at least one excellent peptide match. After the peptide/spectrum matches were filtered, we estimated that the protein FDRs were = 1% for each sample analysis. Resulting protein lists include subset proteins to allow for consideration of all possible protein forms implicated by a given peptide identified from the complex IE protein mixtures. Each protein identified with the IP2 pipeline was associated with several different measures of abundance used in our analyses, including: peptide counts, spectral counts, and normalized spectral abundance factor (NSAF) (Zybailov et al., 2006), which considers protein length and number of proteins identified in the experiment. When comparing ratios and abundances of a given protein across samples, we used NSAF rank rather than abundance to minimize the effects of differences in sample sizes and stochastic differences between MS analyses.

To estimate the number of proteins detected in every experimental condition, we assessed the number of protein entries that displayed detectable NSAF values in the cytosolic, the nuclear or in both subcellular fractions.

To calculate the cytosolic or nuclear ratio of the detected proteins in the different samples, we applied the following equation:

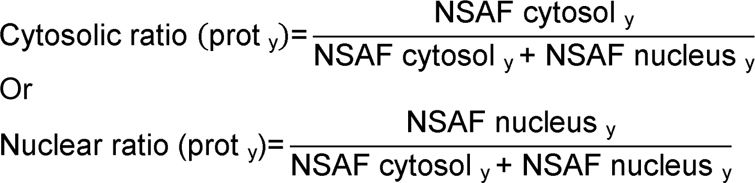

To assess the purity of the subcellular fractionation method utilized in this study, we calculated the cytosolic and nuclear ratio values of *bona fide* nuclear and cytosolic markers in GFP control cells, that were plotted as displayed in Figure 1C. To assess proteomic changes among the multiple transfected conditions tested in this study, we calculated the average ratio (± SD) of both nuclear and cytosolic fractions of every detected protein obtained from 3-4 independent experimental replicates (see Table S1). Obtained results were plotted in violin plots to graphically show the subcellular proteome distribution among the different experimental conditions (Figures 1E, G). We alternatively calculated subcellular ratios utilizing emPAI (exponentially modified protein abundance index) values, which represents the number of observed peptides divided by the number of observable peptides per protein (Figure S1L). Similar results were obtained with both relative abundance factors (NSAF and emPAI).

The comparison between the changes elicited by the expression of C9×58-GFP relative to the ones elicited by the expression of the two control conditions (GFP and C9×8-GFP) was carried out in consultation with the statistics core facility at Northwestern University. We first calculated the average (± SD) of the cytosolic ratio for every detected protein obtained from 4 independent experimental replicates (see Table S1). As shown above, and in Figure 1A, the cytosolic ratio is calculated as the abundance (NSAF) in the cytosolic fraction over the total abundance in both the cytosolic and nuclear fractions. Thus, all values fall between 0-1 and exhibit a non-normal distribution. We compared the cytosolic ratio value for each protein in the (G_4_C_2_)x58-expressing cells *versus* the corresponding value in both control conditions to calculate the magnitude of the shift in the localization for each protein (Δ, delta). In order to ascertain biological significance to these changes and in accordance with other proteome-wide MS-based experiments (Albaum et al., 2011; Fukuda et al., 2017; Poliakov et al., 2011; Tarasova et al., 2017; Zhang et al., 2019), we performed comprehensive non-parametric statistical testing (Kruskal -Wallis rank test, p value < 0.05), commonly used for datasets that do not exhibit a normal distribution. The volcano plot presented in Figure 2A represents two metrics, i.e. -log10 p-value on the y-axis and magnitude of the shift (Δ, delta) on the x-axis, of the changes observed in (G_4_C_2_)x58-expressing cells relative to controls.

Additionally, we considered both of these metrics, as well as broader biological context, to select candidate proteins for further investigation as illustrated in Figure 3A, where we represented the cytosolic (x-axis) and nuclear (y-axis) ratio differences of the proteins that show significant N/C ratio changes (Kruskal -Wallis rank test, p value < 0.05), between (G_4_C_2_)x58-expressing cells and the two control conditions,.

Gene Ontology (GO) analysis was performed using the online tools STRING, PANTHER and DAVID. The COMPARTMENTS (compartments.jensenlab.org) tool was utilized to characterize the normal protein localization patterns of the N/C perturbed proteins and assess the physiological relevance of the observed changes. Bioinformatics & Evolutionary Genomics (ioVe://bioinformatics.psb.ugent.be/webtools/Venn/) and BioVenn, are web applications utilized for the comparison and visualization of biological lists using area-proportional Venn diagrams (Hulsen et al., 2008). NLS sequences detection in the proteins of interest was performed with cNLS Mapper (http://nls-mapper.iab.keio.ac.jp/cgi-bin/NLS_Mapper_form.cgi) and NES sequences with NetNES 1.1 Server (http://www.cbs.dtu.dk/services/NetNES/). To identify NLS and NES sequence motifs in large proteomic datasets, we utilized the NLSdb database (https://rostlab.org/services/nlsdb/).

#### Cell viability analysis

Three different approaches were utilized to assess cell viability in HEK-293 cells transfected with GFP, C9×8-GFP and C9×58-GFP. Firstly, Propidium Iodide (PI) incorporation by apoptotic cells was measured in cells 48hrs after transfection. Cells were dissociated with trypsin, resuspended in DMEM without phenol red (Gibco) + 2% FBS. PI (5µg/ml) was added to the cell suspension right before flow cytometry analysis on BD LSR Fortessa. Secondly, GFP, C9×8-GFP and C9×58-GFP-transfected cells were pulsed with PI for 30min and imaged in a Leica DMi8 microscope to detect double GFP+PI+ cells. Double labeled cells were counted with Fiji software. Thirdly, a colorimetric MTT assay with cells exposed to 3-(4,5-dimethylthiazol-2-yl)-2,5-diphenyltetrazolium bromide (MTT, 0.5mg/ml), which is reduced to formazan (purple) depending on cellular NAPDH activity. Formazan levels were measured by a Synergy HTX Multi-Mode Reader (BioTek) at 500-600nm.

#### Western blot and dot blot analysis

Cells were harvested in RIPA buffer (10mM Tris-HCl pH 8.0, 140mM NaCl, 1mM EDTA, 0.1% SDS), supplemented with 1% Triton X-100 (Triton, Sigma Aldrich), 1mM phenylmethylsulphonyl fluoride (DOT Scientific), and protease inhibitors (Millipore). For western blot analysis lysates were sonicated and protein extracts were separated by SDS-PAGE followed by electrotransfer to a nitrocellulose membrane (Bio-Rad). For dot-blot analysis, 5-10μg of protein was loaded through a 96-well Acrylic Dot-Blot System (Whatman, GE Healthcare Life Sciences) directly onto a nitrocellulose membrane. The membranes were blocked in Tris-buffered saline (TBS, 50mM Tris, 150mM NaCl, HCl to pH 7.6)) + 0.1% Tween 20 (Bio-Rad) + 5% non-fat dry milk (Labscientific) and then incubated overnight at 4°C with primary antibodies: ATF4 (rabbit, 1:500, Cell Signaling Technology), α-tubulin (mouse, 1:5000, Santa Cruz Biotechnology), eRF1 (mouse, 1:500, Santa Cruz), GAPDH (rabbit, 1:1000, Cell Signaling), GFP (goat, 1:1000, Abcam), poly-GP (mouse, 1:1000, generous gift from Tania Gendron), poly-GR (rat, 1:2000, Millipore), GRP78/BiP (mouse, 1:500, Santa Cruz Biotechnology), eIF2α (rabbit,1:1000, Cell Signaling Technology), Phospho-eIF2α (Ser51; rabbit, 1: α Cell α 1000, Signaling Technology), LAMIN A/C (mouse, 1:10000, generously provided by Robert Goldman’s lab), LMNB1 (rabbit, 1:1000, Proteintech), SMG1 (mouse, 1:500, Santa Cruz Biotechnology), TAFII p130 (mouse, 1:1000, Santa Cruz Biotechnology), VINCULIN (mouse, 1:500, Sigma-Aldrich). Primary antibodies were diluted in TBS + 0.1% Tween + 5% BSA (Calbiochem). After several washes in TBS + 0.1% Tween, membranes were incubated with their corresponding secondary HRP-conjugated antibodies (1:5000, LI-COR Biotechnology). Protein signals were detected by a ChemiDoc^TM^ XRS+ (Bio-Rad), using the SuperSignal West Pico chemiluminescent system (Thermo Scientific). Densitometry analysis for the bands was performed using Fiji software (ImageJ, NIH Image).

#### *Drosophila* stocks, maintenance and phenotyping

To identify genetic modifiers of C9-HRE toxicity, *Drosophila melanogaster* lines expressing RNAi for homologues of specific targets were obtained from the Vienna Drosophila Resource Center (eRF1, stock no # 33569, SF2 # 27775, Art1 # 110391; e(y)2 # 108212, Tnpo-SR #40991, CCT8 # 103905) or Bloomington Drosophila Stock Center (Upf1 # 64519) and crossed with a C9ORF72–UAS-(G_4_C_2_)x3 and C9ORF72– UAS-(G_4_C_2_)x30 lines, which were a kind gift from Dr Peng Jin, or a C9ORF72-UAS-(G_4_C_2_)x36 line (Bloomington Drosophila Stock Center). Human eRF1 (hETF1) cDNA was cloned in the pUAST-attb vector by using the VectorBuilder platform. The injection and the generation of transgenic flies was done by Best gene Drosophila Embryo Injection Services by using the site-specific integration of transgene with purified pUAST-attb-hETF1 vector. The UPF1 OE (stock number#33539) and RNAi lines (stock number#64519) were obtained from Bloomington stock center. *Drosophila* stocks utilized in this study were maintained on standard cornmeal medium at 25°C or 18°C in light/dark-controlled incubators. External eye degeneration was used as a screening endpoint as described previously (Lanson et al., 2011; Pandey et al., 2007). Briefly, the following parameters were scored as 0 if normal or 1 if abnormal: level of discoloration, ommatidial disorganization, absence of hairs, presence of necrotic tissue, level of development, shape, and contour. The sum of the scores defined the level of eye degeneration. All data was collected and scored in a blinded manner.

The Rapid Iterative Negative Geotaxis (RING) was performed as described in (Nichols et al., 2012). Briefly, RNAi flies with or without C9-HRE expression in neurons (ELAV-gene switch) were placed into polystyrene vials within the RING apparatus, wherein flies can be pulled to the bottom of vials and climbing movement recorded. Movement was recorded using an iPhone 8 camera, climbing velocity per fly was calculated.

#### Eye sectioning and imaging protocol

Heads of adult female flies (4-day old) were fixed in the Davidson’s fixative, paraffin embedded and sectioned for internal retinal morphology (Goodman et al., 2019). The retinal sectioning and H&E staining were done by Excalibur pathology inc. Images were taken in bright field at 20x in a Leica microscope.

#### *Drosophila* dot blot, Western blot and RT-qPCR analysis

Dot blot was performed as described previously (Mizielinska et al., 2014) with some modifications. Briefly, 4-day old flies were heat-shocked for 1.5 hours at 37°C, snap-frozen and homogenized in RIPA buffer (3μl per fly head) with complete protease inhibitors cocktail (Roche). After adding 6x mli sample buffer (0.5 μl per head), the Laem lysate was heated to 95°C for 5min and 2μl of the samples was blotted onto a dry nitrocellulose membrane. The blot was left to dry for 15 minutes at room temperature and blocked for 1h in 5% BLOT-QuickBlocker reagent in TBS + 0.05% Tween-20 (TBS-T). The membrane was probed with rabbit anti-poly-GR antibody (1:2000 in TBS-T), or mouse anti-tubulin antibody (1:10,000) overnight at 4°C. The following day, the membrane was washed twice with TBS-T and incubated in secondary antibody (anti-rabbit IRDye 800 and anti-mouse IRDye 680D, LICOR, 1: 10,000) for 1h and imaged on Licor imager (Odyssey CLx).

For the eRF1 WB, three adult fly heads were collected from each cross and crushed on dry ice. The granulated heads were incubated in lysis buffer containing 150mM NaCl, 0.1% SDS, 1% NP40, 50mM NaF, 2mM EDTA, 1% sodium deoxycholate, 1mM DTT, 0.2mM sodium orthovandate and protease inhibitor (Roche) for 10min. The lysate was sonicated, centrifuged at 10000xg for 10min and boiled at 95°C for 5min in laemmli buffer. Proteins were separated in a 4-12% NuPAGE gel (Novex/Life technologies) and transferred to a nitrocellulose membrane (Invitrogen). The blot was blocked in 2.5% milk in TBS-T and probed with anti-eRF1 (Santa Cruz Biotechnology) and β-actin (Cell signaling) antibodies overnight at 4°C. The blot was washed in 0.1% TBS-T and incubated with Alexa Flour-tagged secondary for 1h. The blot was imaged in an Odyssey CLx (Li-COR Biosciences).

For RT-qPCR analysis RNA was extracted from fly heads in triplicate (3 heads per replicate) using Trizol (Ambion) and 1-Bromo-3-chloropropane (SIGMA) extraction (Jensen et al., 2013). RNA samples were cleaned using DNA-free kit (Invitrogen) and quantified using a NanoDrop ND-1000 spectrophotometer. cDNA was synthesized from 0.4µg of DNase-free RNA by using the iScript Select cDNA Synthesis Kit (BioRad). qPCR was performed on an Applied Biosystems, 7300 Real Time PCR System by using the iQ Supermix (BioRad) and α ubulin was used as a normalizing control. The comparative Ct method was used for analyzing the results, as previously described (Schmittgen and Livak, 2008), using GraphPad Prism 6 software for statistical analyses.

Primers used for qPCR:

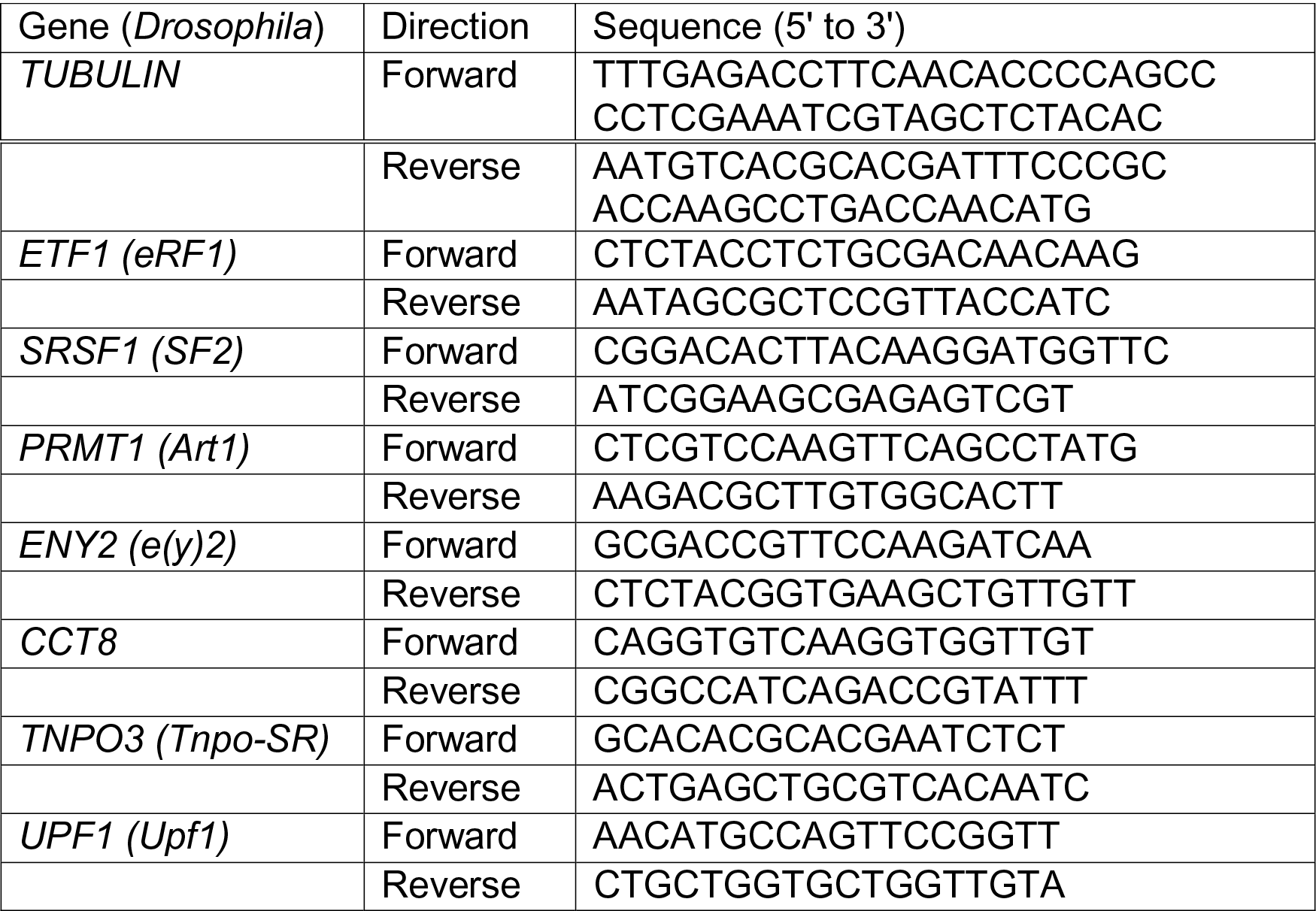

### ELISA

Protein was isolated in the same manner as above. Polyclonal anti-GP capture and detection antibodies in conjunction with Meso Scale Discovery (MSD) electrochemiluminescence detection were used as previously described (Su et al., 2014).

#### RNA preparation and RT-qPCR analysis

RNA was isolated using RNeasy Mini Kit (Qiagen). Briefly, cells were harvested in lysis buffer and gDNA was removed by passing the lysate through a gDNA eliminator column. RNA was then isolated according to manufacturer guidelines and 0.5 to 1µg of RNA was used to generate cDNA using iSCRIPT Reverse Transcription Supermix (Bio-Rad). Real-time qPCR reactions were prepared according to manufacturer guidelines for iTaq Universal SYBR Green Supermix (Bio-Rad) on the CFX system (Bio-Rad). All qPCR assays were performed in triplicate. The average cycle of threshold (Ct) value of two housekeeping genes (ACTIN/GAPDH) was subtracted from the Ct value of the gene of interest to obtain the ΔCt. Relative gene expression was determined as ^2−^ΔCt (ΔΔCt) and expressed relati he control sample or the highest expressed sample experiment.

Primers used for qPCR:

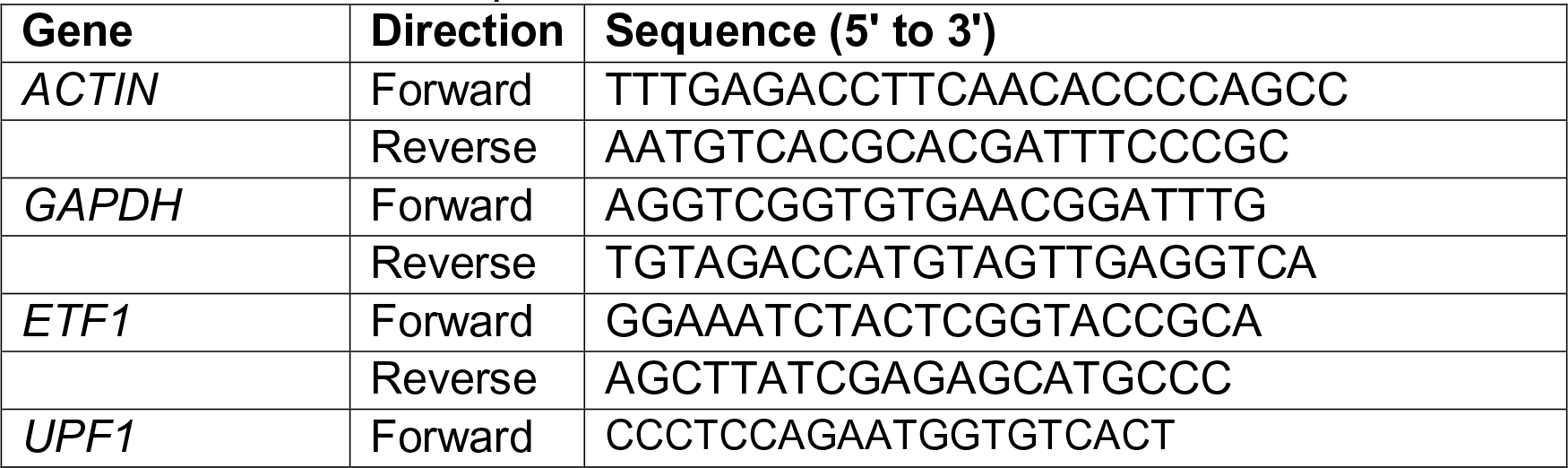
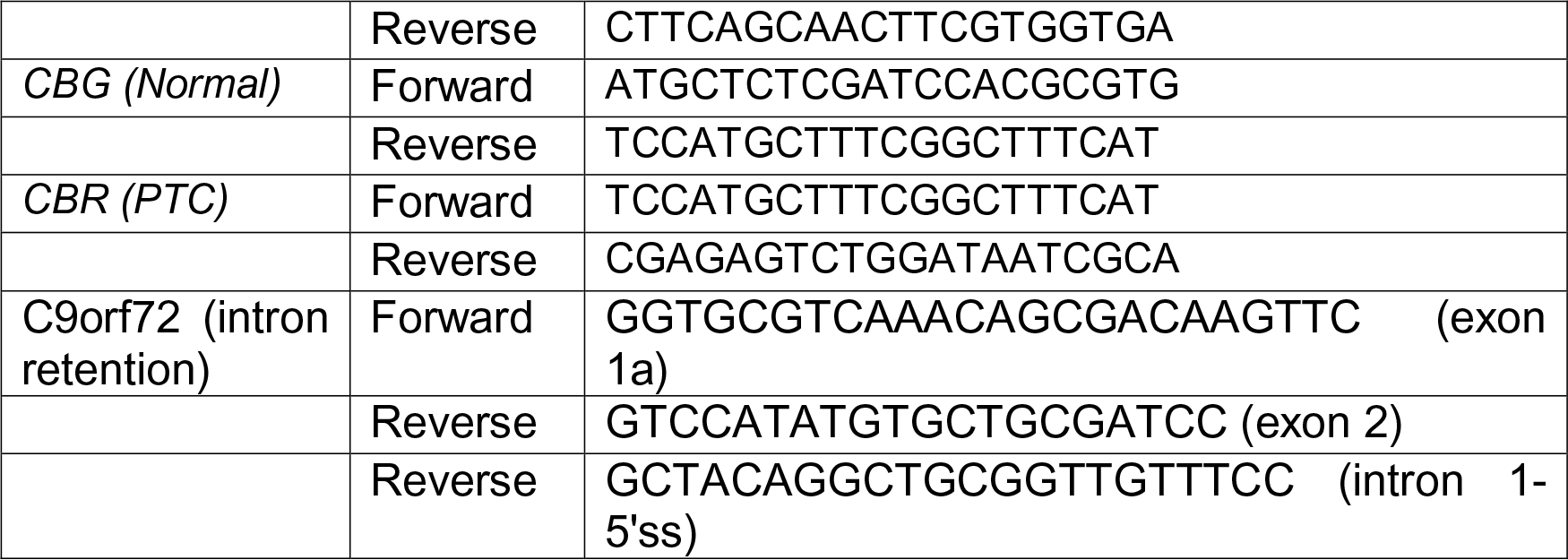

#### Immunocytochemistry

Cells were fixed with 4% paraformaldehyde for 20min, washed with PBS and permeabilized/blocked for 1h in PBS containing 10% normal donkey serum (Jackson ImmunoResearch) and 0.2% Triton. Samples were then incubated overnight at 4°C with primary antibodies: eRF1 (mouse, 1:100, Santa Cruz Biotechnology), GFP (goat, 1:500, Abcam), LAMIN-B1 (rabbit, 1:500, Proteintech), MAP2 (chicken, 1:5000, Abcam), pUPF1-Ser1127 (rabbit, 1:500, Millipore), UPF1 (RENT1, mouse, 1:200, Santa Cruz Biotechnology), ISLET1/2 (mouse, 1:200, DHSB), Puromycin (mouse, 1:10000, Millipore). Next day, PBS + 0.1% Triton was applied for several washes. Samples were then incubated with the appropriate secondary antibodies conjugated to Alexa488, Alexa555 or Alexa647 fluorophores (1:500 to 1:1000 Molecular Probes) for 1h at RT. Cell nuclei were labeled by DNA staining using Hoechst (Life Technologies). Immunolabeled samples were blinded upon mounting for subsequent imaging.

#### Expansion microscopy

Expansion microscopy was performed as previously described (Asano et al., 2018) with minor modifications. In brief, cells were cultured on 12mm coverslips, fixed in 4% PFA, and permeabilized/blocked for 1h in PBS containing 10% NDS and 0.2% Triton. Samples were then incubated overnight at 4°C with primary antibodies: eRF1 (mouse, 1:50, Santa Cruz Biotechnology), LAMIN-B1 (rabbit, 1:100, Proteintech), KDEL (mouse, 1:50, Enzo Life Sciences), KDEL (rabbit, 1:100, Abcam), pUPF1-Ser1127 (rabbit, 1:100, Millipore), Puromycin (mouse, 1:100, Millipore). Next, coverslips were washed with PBS + 0.1% Triton 3 times for 5 min, and then incubated with the appropriate secondary antibodies conjugated to Alexa488, Alexa555 or Alexa647 fluorophores (1:250 Molecular Probes) for 1h at RT. Secondary antibody was removed and PBS + 0.1% Triton was applied 3 times for 5min. Samples were incubated with Hoechst (1:100, Life Technologies) for 10min. Samples were washed with PBS + 10% NDS 3 times for 5min.

Next, samples were incubated in 0.1mg/mL Acryloyl-X, SE (Life Technologies) overnight at RT. Samples were washed with PBS 2 times for 15min. For gelation, perfusion chamber gasket (Invitrogen) was applied to glass slide, and one 12mm coverslip sample was placed on slide inside gasket; 8μL TEMED (Thermo Scientific) and 8μL Ammonium persulfate (Thermo Scientific) were added to 376μL monomer solution (consisting of 9.1% sodium acrylate (Sigma), 2.66% acrylamide (Sigma), 0.16% N,N’-Methylenebis(acrylamide) (Sigma Aldrich), 2.13M NaCl (Ambion), and PBS 1x (Ambion)) and immediately applied to cells. Parafilm-coated glass slide was pressed firmly on gasket. This was repeated for each sample and each gelation assembly was incubated at 37°C for 1h. For digestion, parafilm-coated slide was gently removed, ExM digestion buffer (consisting of 50mM Tris pH 8 (Ambion), 1mM EDTA (Ambion), 0.5% Triton, 800mM Guanidine HCl (Sigma Aldrich), 1% Proteinase K (Qiagen), and PBS 1x) was applied to each sample, and cells incubated for 6hrs at RT. Next, a gel knife was used to transfer each gel into a 100mm petri dish. Water was added to each dish and fresh water was added every hour for 5hrs. Using a gel knife, each gel was divided into 35mm glass-bottom dishes (MatTek). Low-melt agarose (2%, Sigma Aldrich) was applied to immobilize gel on a dish. Images were acquired using a Nikon Plan Apo IR 60x NA 1.27 water immersion objective at matched laser settings and images generated for comparison were processed using identical settings.

#### Immunohistochemistry

Postmortem tissue samples from three ALS patients and three non-neurological controls were obtained through the Northwestern University ALS Clinic and examined by immunohistochemistry. Immunohistochemistry was performed using previously described methods (Deng et al., 2011). Briefly, 6μm sections were cut from formalin fixed, paraffin embedded brain regions containi motor cortex. Sections were µ ng deparaffinized and rehydrated by passing slides in serial solutions: 3 times for 10min each in xylene, 3 times for 5min each in 100% ethanol, 3 times for 3min each in 95% ethanol, once for 5min each in 75% ethanol, 50% ethanol, deionized water, and PBS. Antigen retrieval was performed using a decloaking chamber with a 1X Antigen Decloaker solution (Biocare Medical) at 125°C for 10min. Sections were cooled to room temperature for 30min and rinsed with deionized water. Nonspecific background was blocked with 1% BSA in PBS for 20min at RT. Slides were incubated overnight at 4°C with primary antibodies, rinsed 3 times for 5min with PBS, and then appropriate secondary antibodies conjugated to Alexa488, Alexa555 or Alexa647 fluorophores (1:250, Molecular Probes) diluted in PBS, were applied to slides and incubated at room temperature for 45min. Slides were rinsed 3 times for 5min. To diminish autofluorescence, 0.3% sudan black in 70% ethanol was applied to each section for 45sec, and rinsed in deionized water for 5min. Slides were mounted in ProLong Diamond Antifade Mountant with DAPI (Thermo Scientific) and labels were blinded for subsequent image analysis.

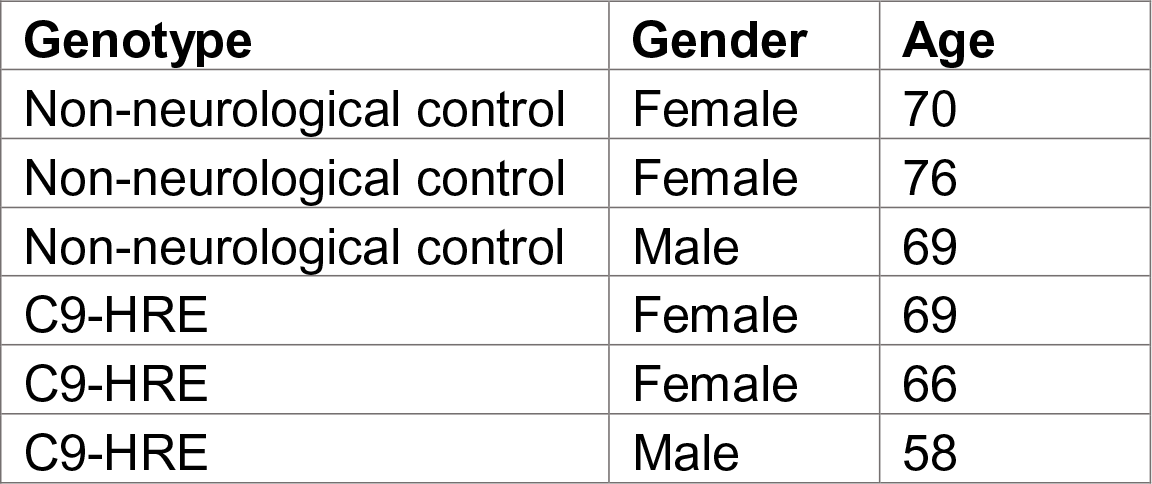

#### *De novo* protein translation analysis

For single-cell protein translation analysis, we utilized the Click-iT HPG protein synthesis kit (Invitrogen). HPG (L-homopropargylglycine, 25µM) is a methionine analog with an alkyne moiety that is incorporated into newly synthesized proteins. Cells were pulsed for 3h with 25μM HPG in Methionine free DMEM (Gibco) and incorporated HPG was detected by Click chemistry following manufacturer guidelines. In short, cells were fixed in 4% PFA for 15min, washed twice with PBS + 0.25% Triton and blocked with PBS + 3% BSA for 1h at RT. Cells were incubated with primary antibodies (ISL1/2, MAP2) overnight at 4°C, then washed with PBS three times, and incubated with the appropriate secondary antibodies. Cells were washed with PBS + 0.25% triton twice and blocked with PBS + 3% BSA for 30min. Next, the click reaction for detection of HPG incorporation was performed, based on a reaction between an azide and alkyne, wherein the alkyne-modified protein is detected with Alexa Fluor® 594 azide. ISRIB (Sigma Aldrich) treatment (2μM) was applied for 5 days in iPSC-derived MNs to assess the potential effect of the ISR in protein translation.

Alternatively, a puromycin-based method termed SUnSET (Schmidt et al., 2009), was used to label newly synthesized proteins. Cell cultures were pulsed for 5-10min with Puromycin (20μM) at 37°C. Cells were then fixed and ICC was carried out as described above utilizing anti-puromycin primary antibody (Millipore).

#### Fluorescent *in situ* hybridization (FISH)

FISH analysis was performed as previously described (Donnelly et al., 2013). Briefly, cells were cultured on 12mm coverslips in 24 well plates, fixed in 4% PFA, and permeabilized in 0.3% Triton for 15min. Before hybridization, cells equilibrated in 2X SSC + 50% formamide for 10min at 60°C and probe mixture was preheated for 10min at 95°C. For hybridization, 200μL of hybridization buffer and 27μL of probe mixture were added to each coverslip and cells were incubated for 1h at 60°C. Following hybridization, cells were incubated with 2X SSC + 50% formamide for 20min at 65°C, then again with fresh 2X SSC/50% formamide for 15min at 65°C, and finally with 1X SSC + 40% formamide for 10min at 60°C. Cells then underwent a series of washes at RT: 3x quick washes with 1X SSC, 2x 5min washes with 1X SSC, a 5min wash with Tris-buffered saline, and a 5min wash with Tris-glycine buffer. Cells were post-fixed in 3% PFA and then incubated with blocking buffer for 1h at RT. Cells incubated with primary antibodies diluted in immunofluorescence buffer overnight at 4°C and secondary antibodies diluted in immunofluorescence buffer for 20min at RT. After secondary antibody incubation, cells were sequentially washed with immunofluorescence buffer, Tris-buffered saline, Tris-glycine buffer, 2mM MgCl_2_ in PBS, and PBS. Coverslips were mounted with ProLong Gold Antifade mounting media with DAPI (Invitrogen) and blinded.

Hybridization buffer consisted of 40% formamide, BSA (2mg/ml, Roche), 1mM ribonucleoside vanadyl complex (Sigma-Aldrich), 10mM NaPO_4_, and 1X SSC. Probe mixture for detection of C9 RNA foci, consisted of 1μL salmon sperm (10μg/ μL, Thermo Fisher Scientific), 0.5μL *E. coli* tRNA (20 μg μL, Thermo Fisher Scientific), 25μL 80% formamide, and 0.4μL 25µM 5’ digoxigenin (DIG)-labeled CCCCGGCCCCGGCCCC locked nucleic acid probe (Exiqon, batch 620574). Probe mixture for detection of poly (A) RNA, consisted of 1μL salmon sperm (10μg/μL, Thermo Fisher Scientific), 0.5μL *E. coli* tRNA (20 μg μL, Thermo Fisher Scientific), 25μL 80% formamide, and 0.8μL 25µM 3’ digoxigenin (DIG)-labeled polyT(25) locked nucleic acid probe (Exiqon, lot 237566815). Tris-buffered saline comprised of 50mM Tris and 15mM NaCl solution, pH 7.4. Tris-glycine buffer comprised of 0.75% glycine and 200mM Tris solution, pH 7.4. Blocking buffer consisted of 1% normal donkey serum (Jackson Immunoresearch) and 5% heat-shocked BSA in Tris-buffered saline. Immunofluorescence buffer consisted of 2% heat-shocked BSA in Tris-buffered saline. DIG-labeled probe was detected with a fluorescein-conjugated sheep anti-DIG antibody (1:250, Roche). All buffers were made with RNase-free water or PBS.

#### Quantitative image acquisition and analysis

Images used for quantification were acquired at matched exposure times or laser settings and processed using identical settings. Quantifications were normalized within each respective experiment with n ≥ 3 independent experiments unless otherwise specified in figure legends. Ima ^≥^acquisition was performed on a Leica DMI4000B laser scanning confocal microscope (Leica, Buffalo Grove, IL) or a Nikon W1 dual camera spinning disk confocal microscope and a Nikon A1R+ laser scanning confocal microscope (Northwestern University Center for Advanced Microscopy).

Unless otherwise specified, image acquisition was performed through the z-dimension at 0.3 – 0.5 μm intervals and individual planes were projected into maximum intensity images. Super-resolution images were acquired using the Nikon Structured Illumination Super-Resolution Microscope (Northwestern University Center for Advanced Microscopy, NIH Grant 1S10OD016342-01) through the z dimension at 0.3μm intervals to achieve ∼115nm resolution in the X-Y plane in multiple colors.

The mean signal intensity of eRF1, pUPF1, and UPF1 for a region of interest was the mean pixel intensity per μ^2^ as calculated by the NIS-Elements Advanced Research v5 software (Nikon software, Northwestern University Center for Advanced Microscopy).

Calculation of tdTomato cytosolic ratio was performed on images acquired using a Leica DMI4000B laser scanning confocal microscope with a 63X oil immersion μ^2^ objective. The mean pixel intensity per of tdTomato for the cytoplasm and the nucleus was quantified by FIJI (NIH) for the calculation of the cytoplasmic to nuclear (C/N) ratio. Treatment with the nuclear export inhibitor KPT-330 (1μM for 30min) was used to validate the efficacy of this N/C transport reporter system.

For the calculation of eRF1 N/C ratio, analysis was performed on single slices since the use of projected maximum intensity images caused false nuclear eRF1 detection originating from a high cytosolic eRF1 signal. For iPSC-derived MNs, images were acquired using a Plan Apo λ 100x 1.45 NA oil immersion objective. For HEK-293 cells, images were acquired using a Plan Apo λ 1.45 NA oil immersion objective. For post-mortem tissue, images of layer V regions of motor cortex tissue were acquired using a Plan Apo λ 60x 1.45 NA oil immersion objective. Corticospinal MNs were identified based size and morphology. For MNs, we carried out blinded on computational analysis based on an algorithm written in NIS-Elements that identifies the nuclear region based on DNA staining, and the cytoplasmic region based on MAP2 labeling, and generated values for mean pixel intensity per μ^2^ of eRF1 for each region For HEK 293 cells, we analyzed cells expressing similar levels and carried out a GFP blinded analysis identifying the nuclear region based on DNA staining, and the cytoplasmic region based on GFP labeling, and generated values for mean pixel μ^2^ intensity per of eRF1 for each region. The N/C ratio was calculated as the mean eRF1-nuclear signal divided by the mean eRF1-cytosolic signal. Nuclear 3-dimensional reconstructions were performed in Imaris (Bitplane software, Northwestern University Center for Advanced Microscopy).

eRF1-DPR colocalization analysis was performed based on the Pearson Correlation Coefficient (PCC) between eRF1 (red) and GFP, (GR)x50-GFP, or (PR)x50-GFP (green) signal within the nucleus of each transfected cell. PCC was calculated by NIS-Elements software.

Calculation of nuclear area occupied by LMNB1+ invaginations: images were acquired using a Plan Apo  λ 60x 1.45 NA oil immersion objective. LMNB1+ invagination area was normalized to the area of the nucleus for each MN. For iPSC-derived MNs, blinded computational analysis was used to calculate the area occupied by LMNB1 invaginations. An algorithm identified the nucleus (based on DAPI+ signal) and then analyzed LMNB1+ signal that was established above a set threshold, and >0.3 μm inside the perimeter of the nucleus. We then calculated the area of LMNB1 invagination normalized to the area of the nucleus for each MN. For post-mortem tissue, the nucleus and LMNB1+ protrusions into the nucleus were outlined manually in a blinded manner on layer V corticospinal neurons.

Calculation of % eRF1 within LMNB1+ nuclear invaginations: images were acquired using a Plan Apo λ 60x 1.45 NA oil immersion objective. Blinded computational analysis was rmed using an algorithm written in NIS-Elements, which identified LMNB1+ nuclear invaginations as described above (nucleus identification was based on DAPI+ signal, and LMNB1+ signal identification above a set threshold, and >0.3 μm inside the perimeter of the nucleus). Finally, the algorithm identified nuclear eRF1 signal in the identified invaginations, and generated values for the number of eRF1 puncta within invaginations and the total number of nuclear eRF1 puncta.

Cell survival, *de novo* translation, and pUPF1 analyses: For cell survival, and *de novo* protein translation analysis, images were acquired at 20x and 100x respectively using a Leica DMi8 microscope (Leica, Buffalo Grove, IL), photographed using a C10600-ORCA-R2 digital CCD camera (Hamamatsu Photonics, Japan) and processed with Fiji.

NMD activation analysis was based on phosphorylated UPF1 (pUPF1) levels (pUPF1/UPF1 signal). For MNs, confocal images were acquired using a Plan Apo λ 60x oil immersion objective and the pUPF1/UPF1 value of each MN was calculate ed. For HEK-293 cells, 20x fluorescent microscopy images were acquired and pUPF1/UPF1 value of each optical field was calculated.

Quantification of G_4_C_2_ RNA foci: Images were acquired using a Plan Apo λ 60x 1.45 NA oil immersion objective through the z dimension at 0.3μm intervals and 3-dimensional images were analyzed by individual plane for nuclear and cytosolic foci in a blinded manner.

Quantification of N/C poly(A) RNA: analysis was performed on single slices since the use of projected maximum intensity images caused false nuclear poly(A) RNA detection originating from cytosolic poly(A) RNA signal. Images were acquired and analyzed in a blinded manner using a Plan Apo λ 60x 1.45 NA oil immersion objective and the poly(A) RNA N/C value of each MN was calculated.

#### Intron retention assay

RNA was isolated and cDNA generated as previously described. PCR reaction was performed using 50ng cDNA template, 0.5μM *C9ORF72*-exon1a forward primer, 0.15μM *C9ORF72*-intron1 reverse primer, 0.5μM *C9ORF72*-exon2 reverse primer, and Phusion Hot Start II DNA Polymerase (Thermo Scientific) with an annealing temperature of 55°C and 33 cycles. Products were run in 1% ethidium bromide gel and imaged with ChemiDoc^TM^ XRS+ (Bio-Rad). Intensity analysis for the bands was performed using Fiji software (ImageJ, NIH Image).

#### NMD reporter assay

The pBS-TCR(PTC)-TCR(WT) NMD reporter plasmid (Nickless et al., 2014) contains 2 minigenes of a T Cell Receptor (TCR), the first one includes a premature stop codon (PTC) and the second unit contains a wild-type (normal) TCR minigene with >99% sequence homology to the first one. The two genes are controlled by separate cytomegalovirus (CMV) promoters. Cells were transfected with UPF1 siRNA and RNA was extracted 3 days later, as a control of NMD inhibition. RT-qPCR analysis was performed to determine the relative abundance of the two TCR transcripts.

### QUANTIFICATION AND STATISTICAL ANALYSIS

All statistical analyses were done with Prism 7 software (GraphPad Software) and in collaboration with Alfred Rademaker, PhD (Professor, Division of Statistics, Northwestern University). We classified an independent biological replicate as either an independent experimental replicate or an independent iPSC differentiation, performed on different days. Individual values were usually displayed by dots in the graphs, and represent all values measured in the study. The sample size (n) of each specific experiment are provided in the results section and the statistical test performed for each specific experiment is defined in the corresponding figure legend. For each statistical analysis, we first tested whether sample data fit into Gaussian distribution using the D’Agostino-Pearson omnibus normality test. To compare two experimental conditions, either a student’s t test (parametric) or a Mann-Whitney U test (non-parametric) was ≥3 experimental conditions, either a one-way ANOVA followed by a Bonferroni post-hoc test (parametric) or a Kruskal-Wallis rank test followed by a two-stage linear step-up procedure of Benjamini, Krieger and Yekutieli (non-parametric) was performed. Analysis of MS/MS datasets was done by one-way ANOVA (Figures 1D), Kruskal-Wallis and Dunn’s multiple comparisons test (Figures 1E, G and S1L), and Kruskal-Wallis rank test (Figure 2A). To compare data distribution between the group of proteins that became significantly translocated in C9×58-transfected cells and all the proteins identified in HEK-293 cells by MS/MS, we used a Chi-square/Fisher’s exact test (Figures 2C and S2B-D). For imaging experiments where we tested for differences in a continuous variable at single-cell resolution, we accounted for variability between independent biological replicates by normalizing all measurements to the average of the corresponding controls. We then tested each biological replicate for differences in variance by the Brown-Forsythe test. In cases where biological replicates tested negative (Figures 1F; 4C; 7A), data from independent biological replicates were pooled per genotype or corresponding condition and tested for significance. In cases where biological replicates tested positive (Figures S4B,D,K; 6B,C,E; 7E), each independent biological replicate was tested for significant differences separately. In such cases, we deemed a biological change as significant only if all independent biological replicates tested were significant, and we display in the corresponding figure the least significant, i.e. the highest p value from all comparisons.

The exact number of biological and/or technical replicates, the specific statistical test applied, and p values from main figures are:

**Figure 1:**

(C) n=3 independent biological replicates.

(D) n=4 independent biological replicates; ANOVA, p=ns, not significant.

(E) n=4 independent biological replicates; Kruskal-Wallis test, p****<0.0001.

(F) n=3 independent biological replicates; Kruskal-Wallis test, p*<0.05, **<0.01.

(G) n=3 independent biological replicates; Kruskal-Wallis test, p****<0.0001.

**Figure 2:**

(A) n=4 independent biological replicates; proteins displaying Kruskal-Wallis test p values<0.05 when comparing (G_4_C_2_)x58-expressing cells to both controls (GFP- and (G_4_C_2_)x8-expressing cells) were labeled in dark gray.

**Figure 3:**

(A) n=4 independent biological replicates; displayed dots represent proteins with Kruskal-Wallis test p values<0.05 when comparing (G_4_C_2_)x58-expressing cells to both controls (GFP- and (G_4_C_2_)x8-expressing cells.

(C) n ≥12 flies; Mann-Whitney U test, p=ns, not significant; ****<0.0001.

(E) n ≥ flies; Mann-Whitney U test, p****<0.0001.

**Figure 4:**

(C) n=3 independent differentiations; Mann-Whitney U test, p****<0.0001.

**Figure 5:**

(B) Statistical differences between 3 non-neurological age-matched controls and 3 C9-ALS patients determined by Mann-Whitney U test, p****<0.0001.

(D) Statistical differences between 3 non-neurological age-matched controls and 3 C9-ALS patients determined by Mann-Whitney U test, p****<0.0001.

**Figure 6:**

(B) n=3 independent differentiations; Kruskal-Wallis test, p**<0.01.

(C) n=3 independent differentiations of one control and two C9-ALS iPSC lines; Kruskal-Wallis test, p=ns, not significant; ****<0.0001.

(E) n=3 independent differentiations; Mann-Whitney U test, p**<0.01. (G) n=6 independent differentiations; T test, p*<0.05.

(H) n=3 independent differentiations; Mann-Whitney U test, p****<0.0001.

**Figure 7:**

(A) n=2 independent differentiations; Mann-Whitney U test, p*<0.05; **<0.01; ***<0.001. (C) n=3 independent differentiations; bars represent the mean ± SEM; Wilcoxon signed rank, p****<0.0001.

(E) n=2 independent differentiations; Kruskal-Wallis test, p=ns, not significant; **<0.01, ****<0.0001.

(F) Left: Statistical differences between 3 control and 3 C9-ALS iPSC lines in n=8 independent differentiations; Mann-Whitney test, p**<0.01. Middle: Statistical differences in C9-ALS MNs transfected with scrambled (Scr) *vs. ETF1* siRNAs. n=6 independent differentiations; T test, p*<0.05. Right: Statistical differences in C9-ALS MNs transfected with scrambled (Scr) *vs. UPF1* siRNAs. n=3 independent differentiations; T test, p*<0.05.

(H) Left: n=18 flies; Mann-Whitney U test, p=ns, not significant; ****<0.0001. Right: n =18 flies except for C9-(G_4_C_2_)x36 + UPF1-RNAi genotypes, which are mostly lethal (n=2); Mann-Whitney U test, p****<0.0001.

## SUPPLEMENTAL TABLES

**Table S1, related to Figure 1.** LC-MS/MS data acquired in HEK-293 cells expressing GFP, C9×8, C9×58, ATX1-32Q, ATX1-84Q, HTT-25Q, or HTT-72Q. The table includes six sheets with information related to figure 1 panels:

Figure 1C: Cytosolic and nuclear ratios of *bona fide* cytosolic and nuclear markers. Figures D-E: LC-MS/MS raw data obtained from GFP, C9×8-, and C9×58-expressing cells.

Figure 1G: LC-MS/MS raw data obtained from GFP, ATX1-32Q-, ATX1-84Q-, HTT- 25Q-, or HTT-72Q-expressing cells.

Figure 1H: Kruskal-Wallis significance rank test for all the redundant proteins identified in cells transfected with plasmids expressing GFP, C9×8, C9×58, ATX1-32Q, ATX1- 84Q, HTT-25Q, or HTT-72Q. List of all proteins that were significantly altered in each one of the three repeat expansion mutations (C9-ATX-HTT comparison).

**Table S2, related to Figure 2.** Comprehensive analysis of the subcellular proteomic differences detected in C9×58-expressing cells. The table includes seven sheets with information related to figure 2 panels:

Figure 2A: Mean ± SD N/C ratio changes of proteins that are redistributed in cells expressing C9×58.

Figure 2C: Molecular weights of redistributed proteins in C9×58-expressing cells, compared to all the proteins detected HEK-293 cells in this study.

Figures 2D-E: Gene Ontology analysis of the group of redistributed proteins in C9×58-expressing cells.

Figure S2A: Comparative subcellular localization analysis of proteins redistributed in cells expressing C9×58.

Figure S2B: NES and NLS domain analysis in proteins redistributed in C9×58-expressing cells compared to the detected HEK-293 cell proteome.

Figure S2C: List of proteins previously identified as nuclear import, export and bi-directional cargoes.

Figure S2D: List of proteins previously identified as C9-HRE RNA and C9-HRE DPR interactors.

**Supplementary Table 3**, Information related to iPSC lines utilized in the present study and the human tissue utilized for IHC analysis.

---

**Figure S1, related to Figure 1.**
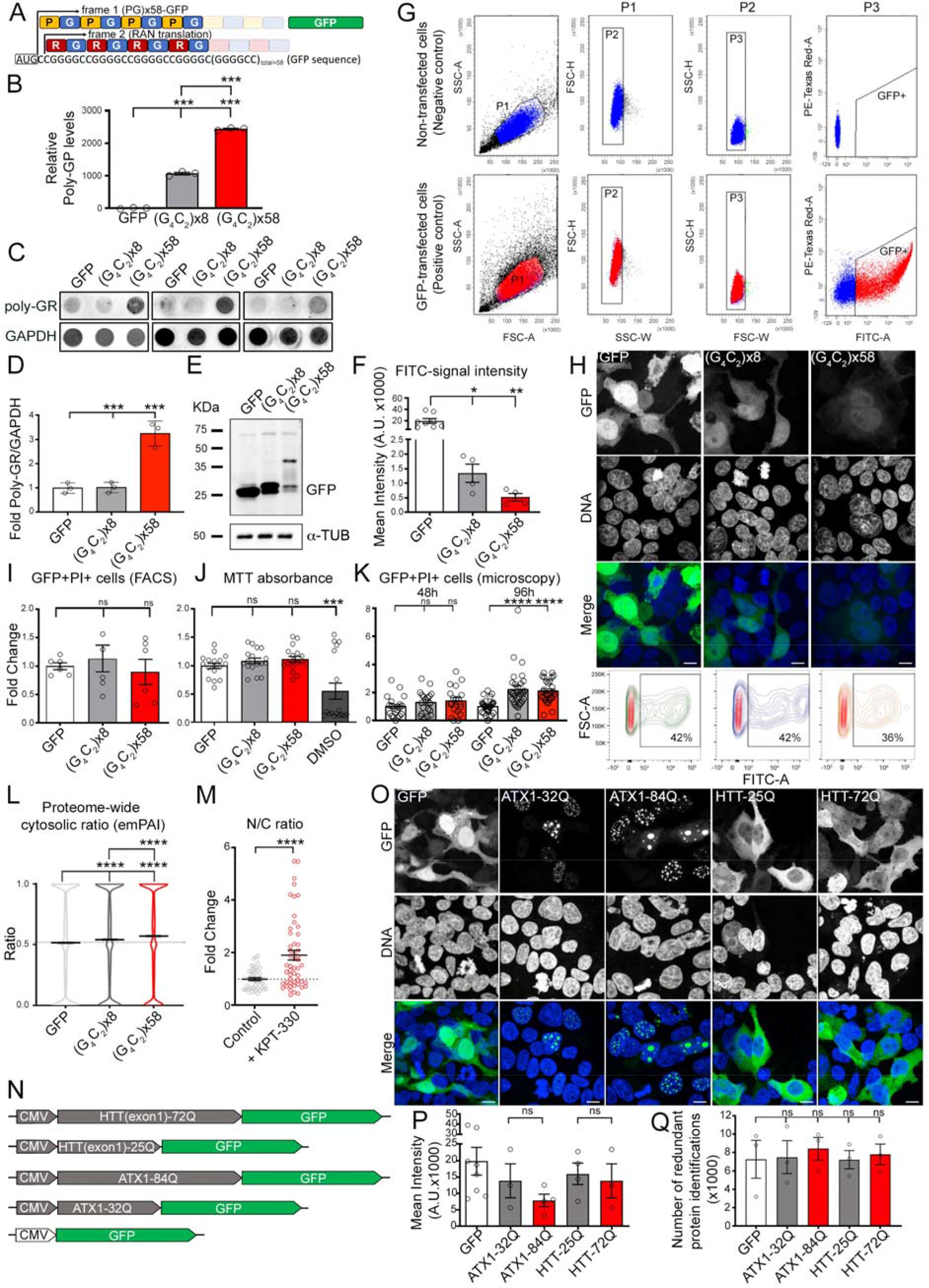
Unbiased Proteomic Analysis of Repeat Expansion Mutations Associated With Neurodegeneration. (A) Schematic showing distinct types of DPRs produced by the C9×58-GFP construct and analyzed in this study. (B) Bar plots representing the average levels of GP DPR detected by ELISA in cells transfected with GFP, C9×8-GFP, or C9×58-GFP. Individual samples are represented as dots (n=3 independent biological replicates; bars represent mean ± SEM; ANOVA, p***<0.001). (C) Dot blot of poly-GR in GFP-, C9×8- or C9×58-expressing cells. (D) Bar plots representing the average levels of poly-GR in cells expressing GFP, C9×8- GFP, and C9×58-GFP. Individual samples are represented as dots (n=3 independent biological replicates; bars represent mean ± SEM; ANOVA, p ***<0.001). (E) Western blot analysis for GFP in HEK-293 cells transfected with GFP, C9×8-GFP or C9×58-GFP. α-TUBULIN was used as a loading control. (F) Bar plots displaying the GFP fluorescence levels acquired by FACS analysis in GFP-, C9×8-GFP- or C9×58-GFP-transfected cells. Individual samples are represented as dots (n=4-7 independent biological replicates; bars represent mean ± SEM; ANOVA, p*<0.05, **<0.01). (G) Representative gating strategy utilized to FACS-purify GFP+ cells. (H) Top: Representative confocal images of HEK-293 cells transfected with GFP, C9×8- GFP, or C9×58-GFP. Scale bars: 20 μm. Bottom: Representative FACS plots depicting effective percentage of sorted GFP+ cells. (I) Bar plots displaying FACS-based quantification of double GFP/Propidium Iodide+ cells from HEK-293 cell cultures 48hrs after transfection with GFP, C9×8-GFP or C9×58-GFP. Individual samples are represented as dots (n=2 independent biological replicates; bars represent mean ± SEM; ANOVA, p=ns, not significant). (J) Bar plots displaying MTT viability assay in HEK-293 cell cultures 48hrs after transfection with GFP, C9×8-GFP or C9×58-GFP. DMSO was used as a positive control. Individual samples are represented as dots (n=3 independent biological replicates; bars represent mean ± SEM; ANOVA, p =ns, not significant, ***<0.001). (K) Bar plots showing percentage of GFP/Propidium Iodide+ cells counted from images of HEK-293 cells 48 and 96hrs after transfection with GFP, C9×8-GFP or C9×58-GFP. Counted fields are displayed as dots (n=3 independent biological replicates; bars represent mean ± SEM; ANOVA, p=ns, not significant, ****>0.0001). (L) Violin plots showing the proteome-wide cytosolic ratios calculated by emPAI values of the ∼10,000 proteins identified in cells expressing GFP (0.516±0.002; mean ± SEM), C9×8-GFP (0.541±0.002), or C9×58-GFP (0.570±0.002) (n=4 independent biological replicates; bars represent mean ± SEM of proteome-wide cytosolic ratio and the dotted line marks the GFP-transfected cells mean value; Kruskal-Wallis test, p****<0.0001). (M) Dot plots displaying the fold change in the N/C ratio of the tdTomato reporter in HEK-293 cells treated with the nuclear export inhibitor KPT-330. Individual cells are represented as dots and the dotted line marks the mean N/C ratio in control samples (n=3 independent biological replicates; bars represent mean ± SEM; Mann-Whitney U test, p****<0.0001). (N) Schematic of the polyglutamine (CAG, polyQ) expansion in *ATX1* or *HTT* (exon 1) used in a comparative subcellular screening of different repeat expansion mutation models. (O) Representative confocal images of HEK-293 cells transfected with GFP, ATX1-32Q, ATX1-84Q, HTT-25Q, or HTT-72Q. Scale bars: 20 μm. (P) Bar plots displaying the mean intensity levels of GFP acquired by FACS analysis in GFP-, ATX1-32Q-, ATX1-84Q-, HTT-25Q-, or HTT-72Q-expressing cells. Individual experimental samples are displayed as dots (n=3-7 independent biological replicates; ANOVA, p=ns, not significant). Q) Bar plots representing the average number of proteins detected in cells expressing GFP, ATX1-32Q, ATX1-84Q, HTT-25Q, or HTT-72Q. Individual experimental samples are displayed as dots (n=3 independent biological replicates; data are represented as mean ± SEM; ANOVA, p=ns, not significant).

**Figure S2, related to Figure 2.**
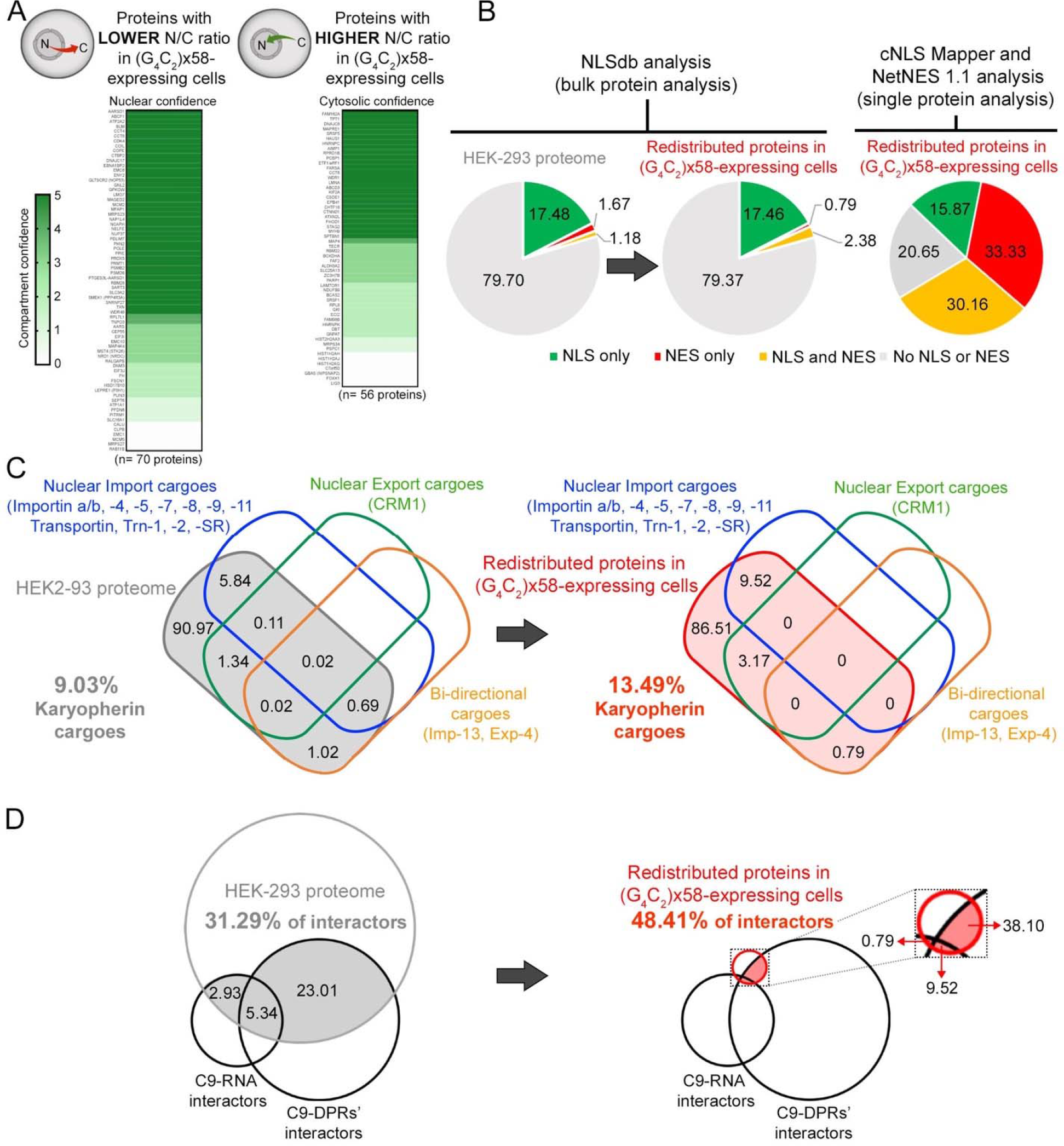
Analysis of Subcellular Proteomic Changes Identified in Cells Carrying the C9-HRE. (A) Heat maps representing the subcellular protein localization analysis of proteins enriched in the cytosol (left) or in the nucleus (right) of C9×58-expressing cells. The presence of these proteins in the nucleus or the cytosol is ranked based on confidence values established by the https://compartments.jensenlab.org database. Color code legend on the left indicates the probability of proteins to be absent (Confidence=0, white) or present (Confidence=1 to 5, from light to dark green) in the nucleus (left) or in the cytosol (right). (B) Pie charts representing the percentage of proteins of the global HEK-293 proteome (left chart) and of the redistributed proteins in C9×58-expressing cells (middle chart) that contain both NLS and NES (yellow), NLS only (green), NES only (red), or neither of these sequences (gray), based on NLSdb analysis. Right pie chart showing the resultant cNLS Mapper and NetNES 1.1 analysis of nuclear localization sequences in the group of redistributed proteins observed in C9×58-expressing cells. (C) Venn diagrams displaying the overlap of previously published nuclear import, export and bi-directional karyopherin cargoes with the HEK-293 proteome (left) or with the group of redistributed proteins in C9×58-expressing cells (right). (D) Venn diagrams displaying the overlap between previously reported C9-HRE-RNA and C9-HRE DPR protein interactors, with the HEK-293 proteome (left) or with the group of redistributed proteins in C9×58-expressing cells (right).

**Figure S3, related to Figure 3.**
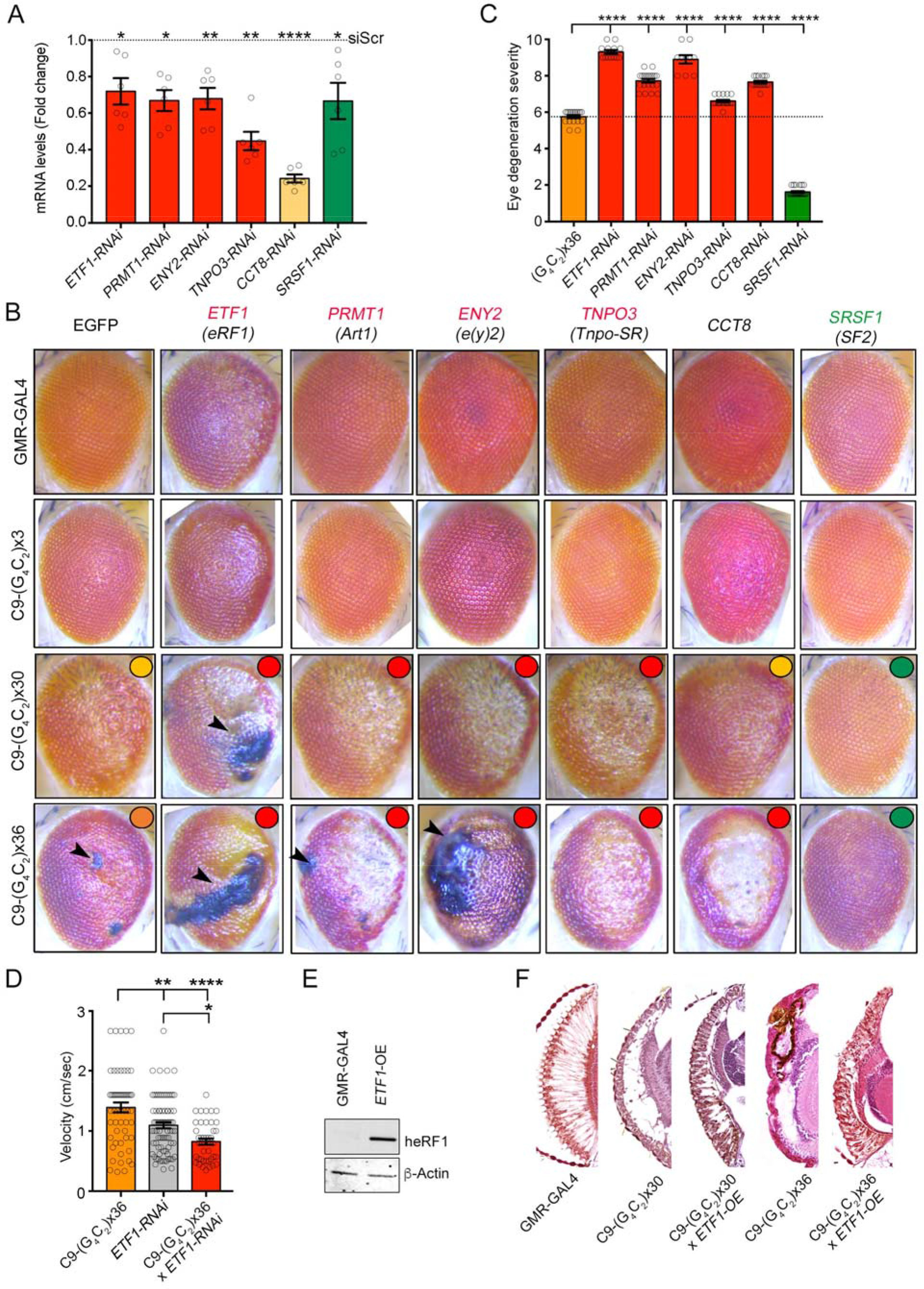
Representative Redistributed Proteins are Genetic Modifiers of *C9orf72* Repeat Expansion Toxicity in *Drosophila* Disease Models. (A) Bar plots displaying RT-qPCR results of target genes in RNAi *Drosophila* models. Individual flies are displayed as dots and dashed line marks the respective mean levels in control flies (n=6 flies; bars represent mean ± SEM; Mann-Whitney U test, p*<0.05, **<0.01, ****<0.0001). (B) Representative eye images of the different C9-ALS flies (C9×30 and x36), and controls (GMR-GAL4 and C9×3) crossed with the multiple knockdown flies tested (*ETF1*, *PRMT1*, *ENY2*, *TNPO3*, *CCT8*, *SRSF1*; homologous gene names in Drosophila are displayed in parenthesis if different). Colored circles on the top right of eye images of C9-ALS flies crossed with the multiple knockdown flies indicate the eye degeneration change on quantifications shown in C: C9×30 toxic level is shown in yellow; C9×36 toxic level is shown in orange; suppression and enhancement of C9-HRE toxicity are shown in green and red, respectively. (C) Bar plots displaying the level of eye degeneration in flies carrying the C9×36 repeats with RNAi for *ETF1*, *PRMT1*, *ENY2*, *TNPO3*, *CCT8*, or *SRSF1*. Enhancers and suppressors of toxicity are represented by red and green bars, respectively. The basal level of eye degeneration in C9×36 flies is shown in orange. Individual flies are represented as dots and the dotted line marks the mean level of eye degeneration in RNAi control C9×36 repeat flies (n ≥ 10 flies; bars represent mean ± SEM; Mann-Whitney U test, p****<0.0001). (D) Bar plots displaying climbing velocity in C9×36, *ETF1*-RNAi and C9×36 x *ETF1*-RNAi flies. Individual flies are displayed as dots (n ≥ 43 flies; bars represent mean ± SEM; ANOVA, p*<0.05, **<0.01, ****<0.0001). (E) Western blot analysis for human eRF1 expression in control (GMR-GAL4) and *ETF1*-OE flies. β-Actin was used as a loading control. (F) Transverse eye sections showing the level of degeneration in C9-HRE flies (C9×30 and x36) with respect to control (GMR-GAL4) and eRF1 overexpression [(C9×30 x *ETF1*-OE) and (C9×36 x *ETF1*-OE)] flies.

**Figure S4, related to Figure 4.**
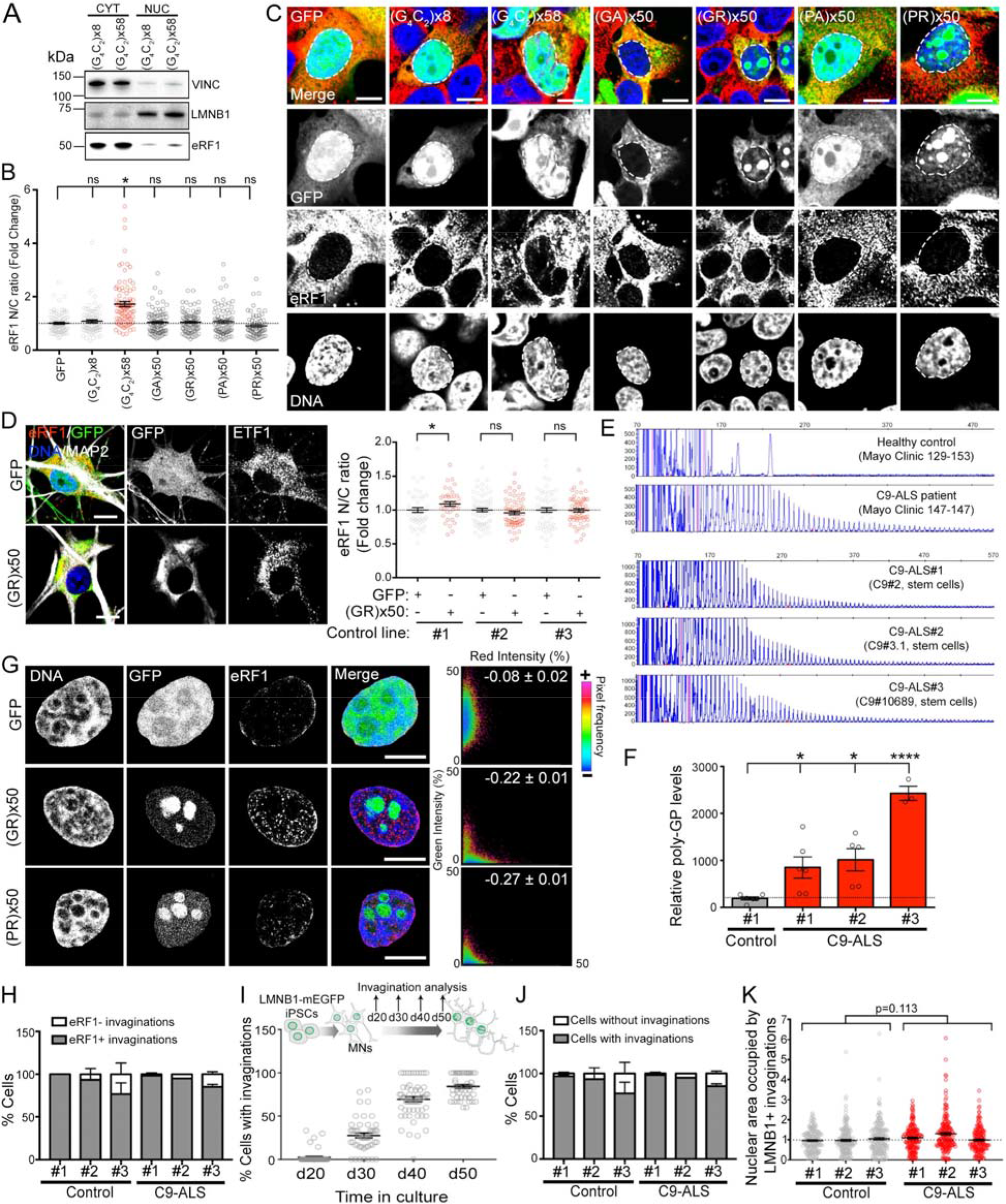
eRF1 is Redistributed in *C9orf72* iPSC-Derived Motor Neurons. (A) Representative western blot bands for eRF1 levels in the nuclear and cytosolic fractions of cells expressing C9×8-GFP and C9×58-GFP. VINCULIN and LMNB1 were used as cytosolic and nuclear markers, respectively. (B) Dot plot displaying the fold change in the N/C ratio of eRF1 signal quantified in HEK- 293 cells expressing GFP, C9×8-, C9×58-, (GA)x50-, (GR)x50-, (PA)x50- or (PR)x50- GFP. Individual cells are displayed as dots and dotted line marks the mean eRF1 N/C ratio in control GFP-expressing cells (n=3 independent biological replicates; bars represent mean ± SEM; Kruskal-Wallis test; p =ns, not significant, *<0.05). (C) Confocal images of GFP, C9×8-, C9×58-, (GA)x50-, (GR)x50-, (PA)x50- or (PR)x50- GFP transfected cells immunolabeled for eRF1 (red) and DNA (blue). Dashed line marks the nuclear boundary. Scale bars: 10 μm. (D) Left: Representative confocal images of healthy control MNs infected with GFP (top panel), or (GR)x50-GFP (bottom panel) and immunolabeled for eRF1 (red), MAP2 (white), and DNA (blue). Scale bars: 10 μm. Right: Dot plot showing the fold change in the N/C ratio of eRF1 signal in MNs derived from 3 healthy control iPSC lines infected with GFP or (GR)x50-GFP. Individual cells are displayed as dots and dotted line marks the mean N/C ratio of eRF1 signal in GFP-infected iPSC-derived MNs (n=2-3 independent differentiations; bars represent mean ± SEM; Mann-Whitney U test, p=ns, not significant, *<0.05). (E) Repeat primed PCR analysis on 3 C9-ALS patient iPSC lines utilized in this study. Healthy control, and C9-ALS patient samples (top) were utilized as a negative and positive control, respectively. (F) Bar plots showing the poly-GP levels detected by ELISA in 1 control and 3 C9-ALS iPSC-derived MNs. Individual samples are represented as dots and the dotted line marks the mean levels of poly-GP in control iPSC-derived MNs (n=3-5 independent differentiations; bars represent the mean ± SEM; ANOVA, p*<0.05; ****<0.0001). (G) Left: Representative confocal images of HEK-293 nuclei expressing GFP, (GR)x50- or (PR)x50-GFP (green), and immunolabeled for eRF1 (red) and DNA (blue). Scale bars: 5 μm. Right: Scatter plots where each point reflects the DPR-GFP (green) signal intensity on the y-axis, and eRF1 signal intensity (red) on the x-axis. The mean ± SEM of the Pearson’s correlation coefficient for the entire dataset is displayed within the corresponding plot. (H) Bar plot showing the percentage of controls and C9-ALS iPSC-derived MNs that contain nuclear invaginations with (gray bars) or without (white bars) eRF1. (I) Top: Schematic illustrating the time course analysis of nuclear invaginations in LMNB1-mEGFP iPSC-derived MNs carried out. Bottom: Dot plot displaying the percentage of LMNB1-mEGFP iPSC-derived MNs containing nuclear invaginations at day 20, 30, 40 and 50 of MN differentiation. (J) Bar plot showing the percentage of controls and C9-ALS iPSC-derived MNs with (gray bars) or without (white bars) nuclear invaginations at day 50 of differentiation. (K) Dot plot displaying the fold change of the nuclear area occupied by nuclear invaginations in MNs derived from 3 controls and 3 C9-ALS patient iPSC lines. Individual cells are represented as dots and the dotted line marks the mean occupied area in control samples (n=3 independent differentiations; bars represent mean ± SEM; Mann-Whitney U test, p=0.113).

**Figure S5, related to Figure 5.**
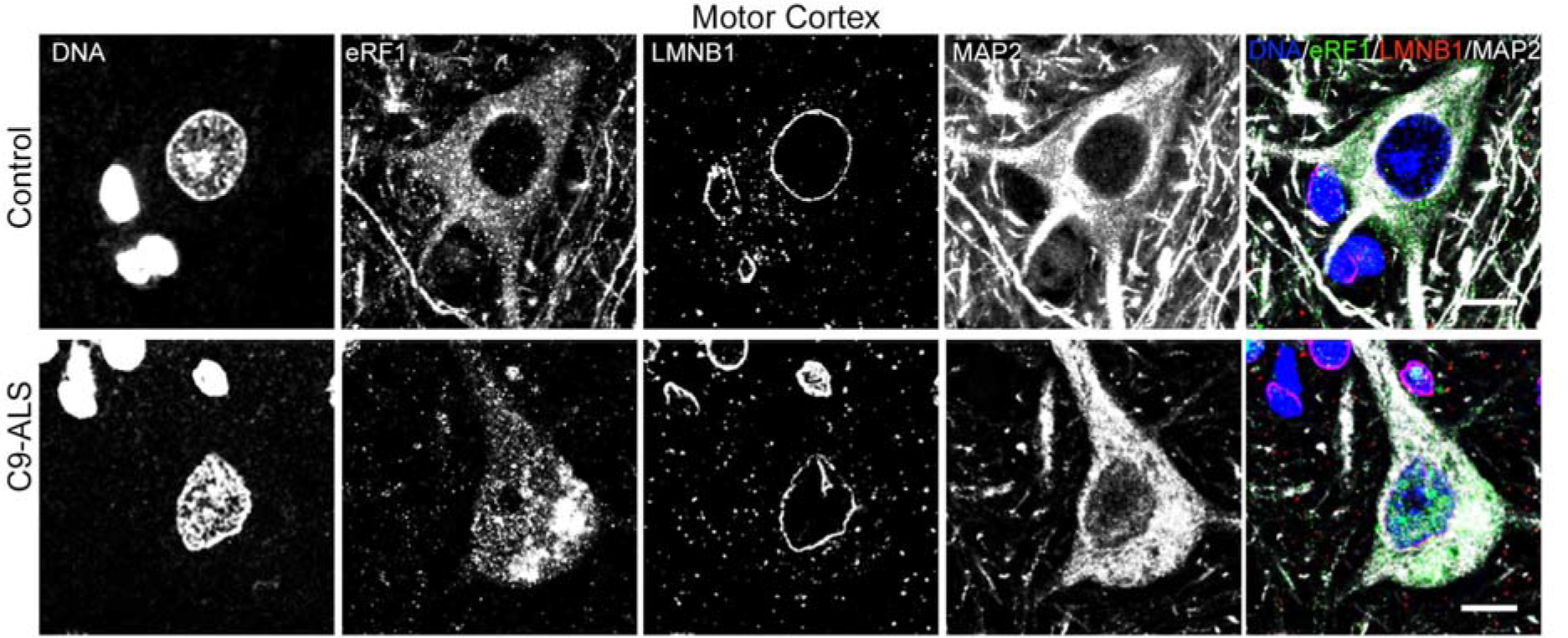
eRF1 Accumulates in Neuronal Nuclei of *C9orf72* ALS Patients Postmortem Tissue. Representative IHC confocal images of layer V neurons immunolabeled for eRF1 (green), LMNB1(red), MAP2 (white), and DNA (blue) in motor cortex tissue from a non-neurological control and a C9-ALS patient. Scale bars: 10μm.

**Figure S6, related to Figure 6.**
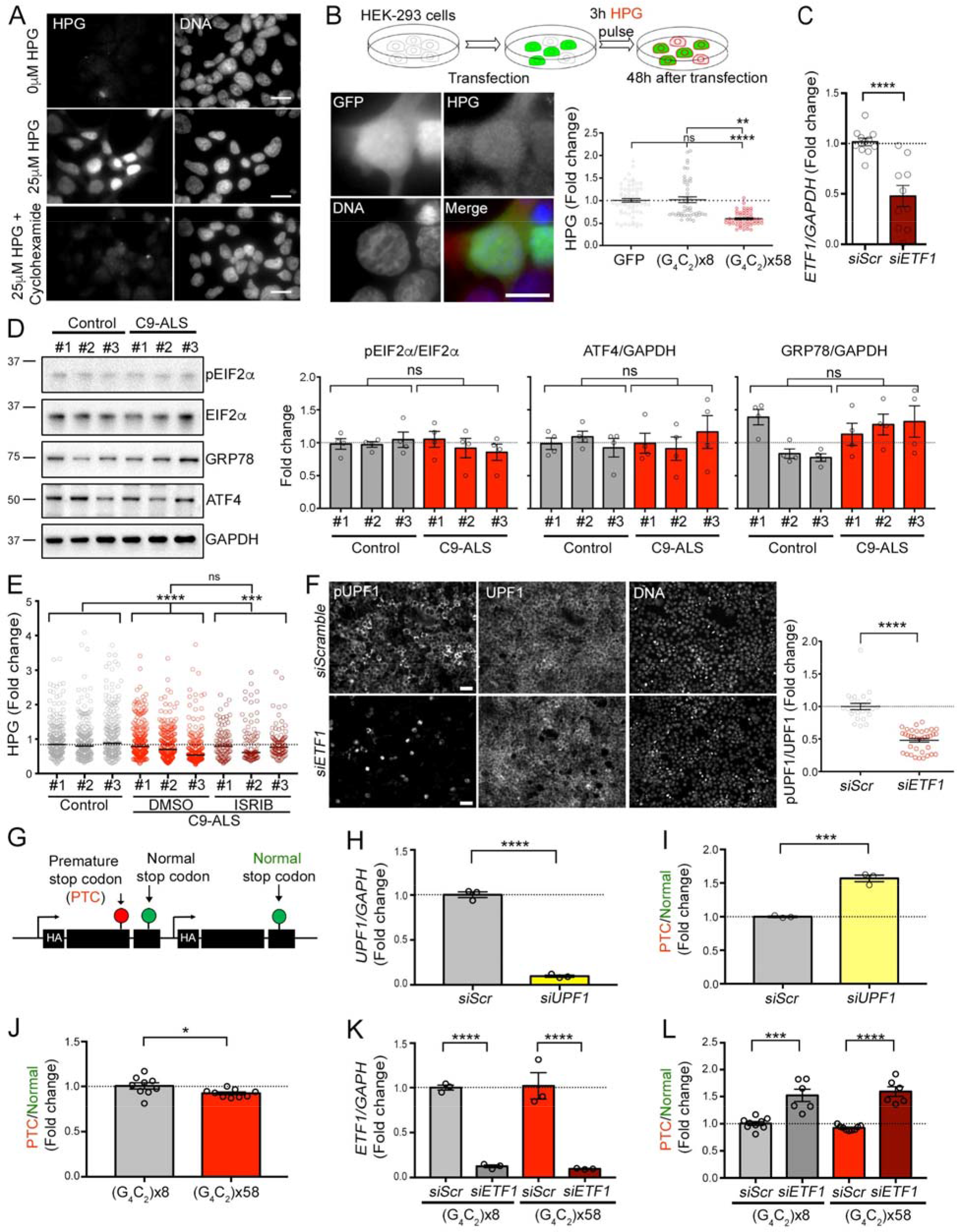
eRF1 is a Key Regulator of Protein Translation and NMD. (A) Quality control analysis of the Click-iT HPG method utilized to measure *de novo* protein translation. Representative images of HEK-293 cells labeled for HPG incorporation and DNA, showing differential fluorescence intensity between cells treated with 25 μM HPG (middle) and untreated cells (top), or cells treated with the translation inhibitor cycloheximide (CHX; bottom). Scale bars: 20μm. (B) Top: Schematic of *de novo* protein translation assay performed in transfected HEK- 293 cells. Bottom left: Representative image of Click-iT HPG fluorescent labeling of *de novo* protein synthesis in GFP-expressing cells. Scale bar: 20 μm. Bottom right: dot plot displaying the levels of *de novo* protein translation assessed by HPG incorporation in GFP-, C9×8- and C9×58-expressing cells. Individual cells are represented as dots and the dotted line marks the mean level in the GFP control condition (n=3 independent biological replicates; bars represent mean ± SEM; Kruskal-Wallis test, p=ns, not significant, **<0.01, ****<0.0001). (C) Dot plot showing the *ETF1* mRNA levels, analyzed by qPCR, in control and C9-ALS iPSC-derived MNs transfected with scrambled or *ETF1* siRNA. *GAPDH* was used as housekeeping gene. Individual samples are represented as dots and the dotted line marks the mean *ETF1* expression levels in siScr-transfected MNs (n=3 independent differentiation; bars represent mean ± SEM; T test, p****<0.0001). (D) Left: Representative western blot analysis for ISR markers (pEIF2α, GRP78 and ATF4) in MN cultures derived from 3 control and 3 C9-ALS iPSC lines. EIF2α and GAPDH were used as loading controls. Right: bar plots displaying the fold change in the protein levels of ISR markers in 3 C9-ALS *vs.* 3 healthy control iPSC-derived MNs. Individual samples are displayed as dots and dotted lines mark the mean ratio value in control samples (n=4 independent differentiations; bars represent the mean ± SEM; T test, p=ns, not significant). (E) Dot plot displaying the fold change of HPG incorporation in MNs derived from 3 controls and 3 C9-ALS patient iPSC lines treated with vehicle (DMSO) or ISR inhibitor (ISRIB). Individual cells are represented as dots and the dotted line marks the mean protein translation in control samples (n=3 independent differentiations; bars represent median; Kruskal-Wallis test; p=ns, not significant, ***<0.001, ****<0.0001). (F) Left: ICC analysis of NMD activation as measured by the pUPF1/UPF1 ratio upon *ETF1* knockdown in HEK-293 cells. Scale bar: 50 μm. Right: Dot plot showing the pUPF1/UPF1 levels in HEK-293 cells transfected with scramble and *ETF1* siRNAs. Individual analyzed fields are displayed as dots and the dotted line marks the ratio mean in the siScr control condition (n=3 biological replicates; bars represent mean ± SEM; Mann-Whitney U test, p****<0.0001). (G) Schematic of the reporter construct utilized to assess NMD activity *in vitro*. The construct encodes two TRB (T cell receptor-β) minigenes sharing >99% homology and fused to an HA tag; one of them contains a premature stop codon (PTC) that produces a truncated protein. (H) Dot plot showing qPCR analysis of *UPF1* mRNA levels in HEK-293 cells transfected with scrambled or *UPF1* siRNA. *GAPDH* was used as housekeeping gene. Individual samples are represented as dots and the dotted line marks the mean *UPF1* expression levels in siScr-transfected cells (n=3 independent biological replicates; bars represent mean ± SEM; T test, p****<0.0001). (I) qPCR analysis of the ratio of truncated (PTC) to full-length (normal) RNA products of the TRB minigenes in HEK-293 cells transfected with scrambled or UPF1 siRNAs. Individual samples are represented as dots and the dashed line marks PTC/Normal ratio in siScr-transfected HEK-293 cells (n=3 independent biological replicates; bars represent mean ± SEM; T test, p***<0.001). (J) Bar plot displaying the quantification of the average NMD activity, calculated by the PTC relative to full-length product in cells expressing C9×8-GFP or C9×58-GFP. Independent replicates are displayed as dots and dotted line marks the mean PTC/normal ratio in control samples (n=3 independent biological replicates; bars represent mean ± SEM; T test, p*<0.05). (K) Dot plot depicting the RT-qPCR analysis of *ETF1* mRNA levels in C9×8- and C9×58-expressing cells co-transfected with scrambled or *ETF1* siRNA. *GAPDH* was used as housekeeping gene. Independent replicates are displayed as dots and dotted line marks the mean *ETF1* levels in siScr-transfected HEK-293 cells (n=3 independent biological replicates; bars represent mean ± SEM; T test, p****<0.0001). (L) Dot plot displaying the quantification of NMD activity by qPCR in HEK-293 cells expressing C9×8-GFP and C9×58-GFP with normal or reduced levels of *ETF1*. Individual samples are displayed as dots and dotted line marks the mean PTC/normal ratio in siScr-C9×8-transfected cells (n=2 biological replicates; horizontal bars represent mean ± SEM; ANOVA, p***<0.001, ****<0.0001).

**Figure S7, related to Figure 7:**
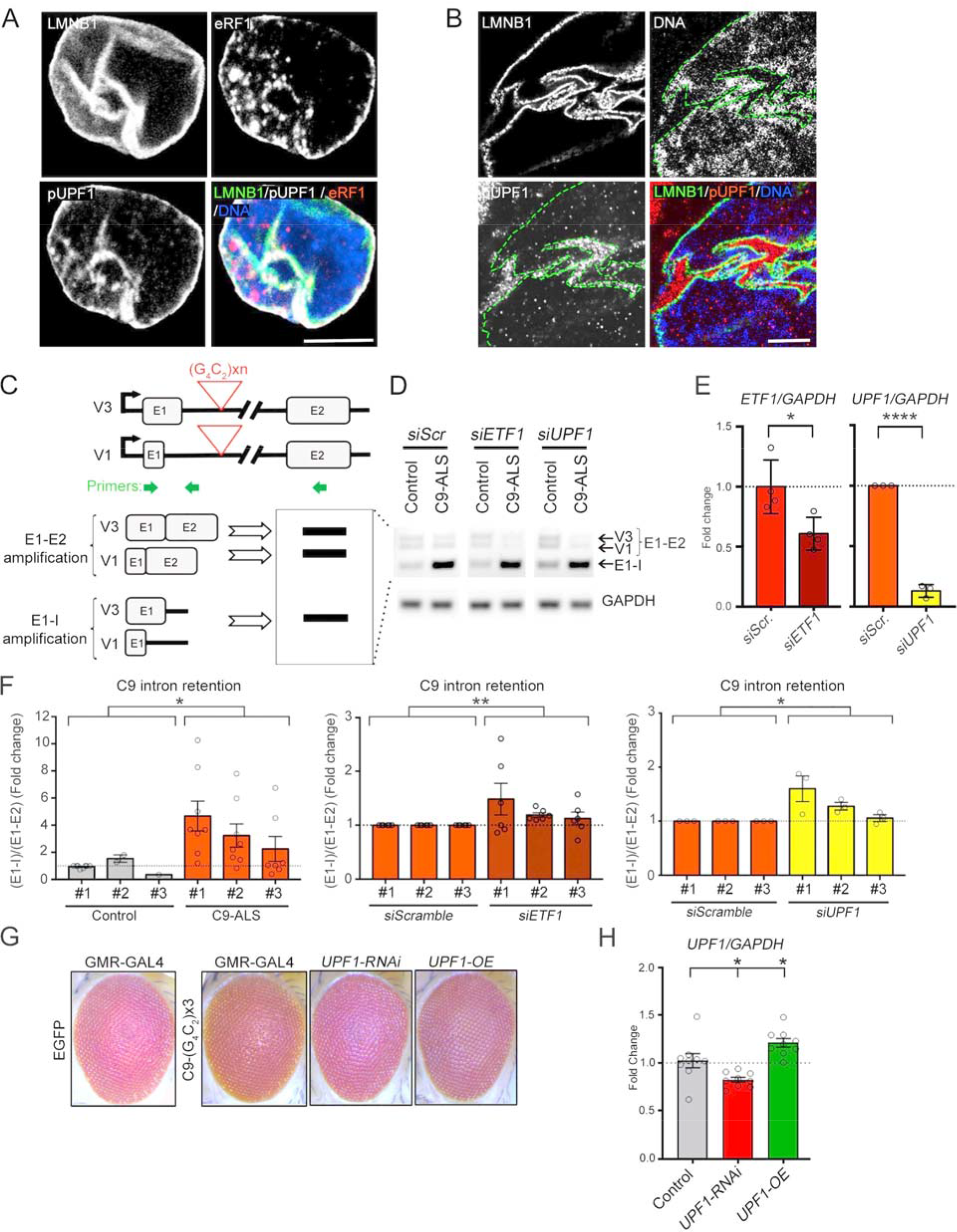
NMD Targets the Expanded *C9orf72* Transcript. (A) Representative confocal image of an iPSC-derived MN nucleus labeled for eRF1 (red), pUPF1 (white), LMNB1 (green), and DNA (blue), showing high degree of colocalization between eRF1 and pUPF1 within LMNB1+ nuclear invaginations. Scale bar: 5 μm. (B) Representative expansion microscopy image of a C9-ALS patient MN nucleus immunolabeled for LMNB1 (green), pUPF1 (red), and DNA (blue), showing the accumulation of pUPF1 (red) on the cytosolic side of nuclear invaginations. Dashed lines indicate the nuclear envelope border. Scale bar: 10 μm. (C) Schematic of the C9 intron retention assay utilized in this study. (D) RT-PCR-based C9 intron retention analysis, by comparing the ratio of the mutant (E1-I) relative to wild type (E1-E2) transcript in 1 control and 3 C9-ALS patient-derived MNs treated with scramble, *ETF1*, or *UPF1* siRNAs. Control and C9-ALS adjacent bands for each transfected condition, from the same gel and acquisition time exposure, were cropped to display representative changes observed in the multiple tested conditions. (E) Bar plots showing qPCR analysis of *ETF1* (left) and UPF1 (right) mRNA levels in C9-ALS iPSC-derived MNs transfected with scrambled and *ETF1* or *UPF1* siRNAs, respectively. *GAPDH* was used as housekeeping gene. Individual samples are represented as dots and the dotted line marks basal levels of *ETF1* and *UPF1* in siScr- C9-ALS iPSC-derived MNs (n=4, left graph, and n=3, right graph independent differentiations; data are represented the mean ± SEM; T test, p*<0.05, ****<0.0001). (F) Bar plots showing fold change levels of C9-intron retention, as measured by RT-PCR, obtained from MN cultures derived from 3 control and 3 C9-ALS iPSC lines (left graph), in C9-ALS iPSC-derived MNs transfected with scrambled (Scr) *vs. ETF1* siRNAs (middle graph), and in C9-ALS iPSC-derived MNs transfected with Scr *vs. UPF1* siRNAs (right graph). Individual samples are displayed as dots and dotted line marks the mean C9 intron retention in controls (left graph) and in siScr-transfected C9- ALS (middle and right graphs) MNs (n=8 (left), 6 (middle), 3 (right) independent differentiations; bars represent the mean ± SEM; Mann-Whitney (left and right graphs), and t test (middle graph), p*<0.05; ***<0.01). (G) Representative images of fly eyes from wild-type (GMR-GAL4 x EGFP) or C9×3 control mutant flies, with endogenous levels (EGFP), knockdown (*UPF1*-RNAi) or overexpression (*UPF1*-OE) of UPF1. (H) Dot plot showing qPCR analysis of *UPF1* mRNA levels in the Drosophila models of *UPF1* knockdown (*UPF1*-RNAi) and overexpression (*UPF1*-OE). *GAPDH* was used as housekeeping gene. Individual flies are represented as dots and the dotted line marks the mean *UPF1* levels in control flies (n=9 flies/condition; bars represent mean ± SEM; ANOVA, p *<0.05).

